# A multiscale theory for spreading and migration of adhesion-reinforced mesenchymal cells

**DOI:** 10.1101/2023.05.03.539193

**Authors:** Wenya Shu, C. Nadir Kaplan

## Abstract

We present a chemomechanical whole-cell theory for the spreading and migration dynamics of mesenchymal cells that can actively reinforce their adhesion to an underlying viscoelastic substrate as a function of its stiffness. Our multiscale model couples the adhesion reinforcement effect at the subcellular scale with the nonlinear mechanics of the nucleus-cytoskeletal network complex at the cellular scale to explain the concurrent monotonic area-stiffness and non-monotonic speed-stiffness relationships observed in experiments: We consider that large cell spreading on stiff substrates flattens the nucleus, increasing the viscous drag force on it. The resulting force balance dictates a reduction in the migration speed on stiff substrates. We also reproduce the experimental influence of the substrate viscosity on the cell spreading area and migration speed by elucidating how the viscosity may either maintain adhesion reinforcement or prevent it depending on the substrate stiffness. Additionally, our model captures the experimental directed migration behavior of the adhesion-reinforced cells along a stiffness gradient, known as durotaxis, as well as up or down a viscosity gradient (viscotaxis or anti-viscotaxis), the cell moving towards an optimal viscosity in either case. Overall, our theory explains the intertwined mechanics of the cell spreading, migration speed and direction in the presence of the molecular adhesion reinforcement mechanism.

## 1 Introduction

Cell motion plays a crucial role in various biological processes ranging from tissue formation to metastasis. The mechanical properties of the extracellular matrix (ECM) are critical environmental cues that orchestrate the spreading and migration of adherent (mesenchymal) cells [1–8]. Focal adhesion (FA) sites play a key role in cell mechanosensitivity by mediating the chemical and mechanical interactions between the cell and ECM. The FA dynamics can directly impact the chemically driven localized protrusions and contractions across the cell, governed by the tightly coupled spatiotemporal distributions of the proteins, such as the Rho GTPases, ROCK, or CDC42, which influence the migration dynamics [9–14]. Thus, a rigorous mathematical description of cell migration must include accurate modeling of the adhesion forces at different FA sites and their coupling with the cytoskeletal dynamics driven by the intracellular signaling pathways.

Although some cells such as the U-251MG glioblastoma cells exhibit a biphasic dependence of the traction force on the rigidity of the underlying matrix [15, 16], which can be explained by the classical motor-clutch theory [17, 18], many other systems, such as endothelial cells and fibroblasts, display a monotonic rigidity-force relationship [19, 20]. This behavior is attributed to active adhesion reinforcement triggered by the unfolding of the talin molecules that leads to the vinculin binding and in turn to an increased integrin density at an FA site [21, 22]. While an augmented motor-clutch theory with adhesion reinforcement successfully captures the monotonic increase of the traction force with the matrix stiffness [23], it cannot explain the non-monotonic dependence of the migration speed on the ECM rigidity measured in the experiments [5–8]. This suggests that the mutual dynamics of the many FA sites across the cell along with the cytoskeletal dynamics must be considered to relate the migration speed to the traction forces and in turn to the matrix rigidity, rather than at a single FA site. To reproduce the cell migration dynamics as a coordinated sequence of peripheral protrusion and contractions [4, 24, 25], whole-cell level formulations with multiple FA sites have been developed, including our recent theory that integrates the chemomechanical coupling between the Rho GTPase concentrations, the FA sites, the cytoskeletal network and the nucleus dynamics [26–28]. However, none of these studies have incorporated the adhesion reinforcement mechanism into the whole-cell dynamics. Importantly, we anticipate that a mere addition of the adhesion reinforcement effect to the whole-cell-scale model would still lead to a monotonic rigidity-speed relationship, implying further subtleties associated with the collective intracellular dynamics under adhesion reinforcement.

Here we generalize our multiscale theory in Ref. [28] to elucidate the spreading and migration dynamics of adhesion-reinforced cells on viscoelastic substrates, overcoming the limitations of our previous model without adhesion reinforcement. Our model demonstrates how the nonlinear mechanics of the nucleus–cytoskeletal network complex must play a critical role in the experimental spreading and non-monotonic stiffness-speed profiles: Large cell spreading on stiff substrates induced by the augmented local traction forces due to adhesion reinforcement, an effect accurately captured by our model, deforms the nucleus [29] and increases both the cytoplasmic viscosity and cytoskeletal stiffness [8, 30–32]. Consequently, the drag force on the nucleus increases, making it harder for the cell to translocate the nucleus on stiff substrates, as evidenced by a theoretical non-monotonic stiffness-speed relationship in quantitative agreement with experiments. By using the standard linear solid model for the substrate viscoelasticity, we also investigate the effect of the substrate stress relaxation on cell migration across a broad stiffness range, which is experimentally well documented [23, 33–35]. We show that, on a substrate with high elastic stiffness, fast stress relaxation (low viscosity) may preempt the effect of adhesion reinforcement whereas slow relaxation (high viscosity) may promote it. That way, our model quantitatively reproduces the experimental spreading area and migration speed profiles of the HT-1080 human fibrosarcoma cells [27]. Furthermore, we demonstrate that adhesion-reinforced cells persistently move up the stiffness gradients, i.e., durotaxis [36–38], in good agreement with the experimental migration patterns of the MDA-MB-231 cells [16]. We also reveal the effect of the viscosity gradients on directed migration, a.k.a. viscotaxis, which sheds light on the migration of human mesenchymal stem cells from high to low loss moduli on collagen coated polyacrylamide gels [39].

Our theory provides a rigorous understanding of how the nucleus drag as a function of cell spreading gives rise to the experimentally observed complex migration dynamics, in contrast with the previous phenomenological treatments that assumed a direct functional relationship between the drag force, the traction force, and substrate stiffness [40, 41]. Furthermore, because our model takes into account additional intracellular components beyond just the FA sites, it provides a plausible mechanism for the emergent nonuniform stiffness-speed relation that complements the cell model which relates cell speed only to the resultant traction force from the collective adhesion-reinforced FA dynamics [42].

## 2 Methods

Mesenchymal migration entails the spatiotemporal coordination of front protrusion driven by actin polymerization and focal adhesion (FA) followed by rear contraction due to focal de-adhesion and actomyosin activity. The front-rear symmetry is primarily broken by the chemical polarization of the active and inactive Rho GTPase proteins, which govern the dynamics of the FA sites.

Here we generalize our previous multiscale framework for the talin-low cell migration, which accounted for this complexity by considering the feedback between the biological components in Fig. 1 A, by including the effect of focal adhesion reinforcement due to talin unfolding (Fig. 1 B) [28]. Our modified multiscale model takes into account the nonlinear strain-stiffening of the cytoskeleton and the augmented viscous drag force on the deformed cell nucleus under cell flattening, as well as uses the standard linear solid (SLS) model for the ECM viscoelasticity. At the subcellular scale, a motor-clutch model with adhesion reinforcement is employed to determine the effect of substrate rigidity on traction forces at each FA site (Fig.1 C). Since cells operate in low Reynolds numbers, we only consider the viscous forces arising from the nucleus-cytoplasm and cell membrane-substrate frictions at the cellular scale. In our model, the forces across the two scales are transmitted by the cytoskeleton, and the augmented viscous drag force on the deformed cell nucleus in turn determines the active motion of the cell (Fig.1D and E).

**Figure 1:**
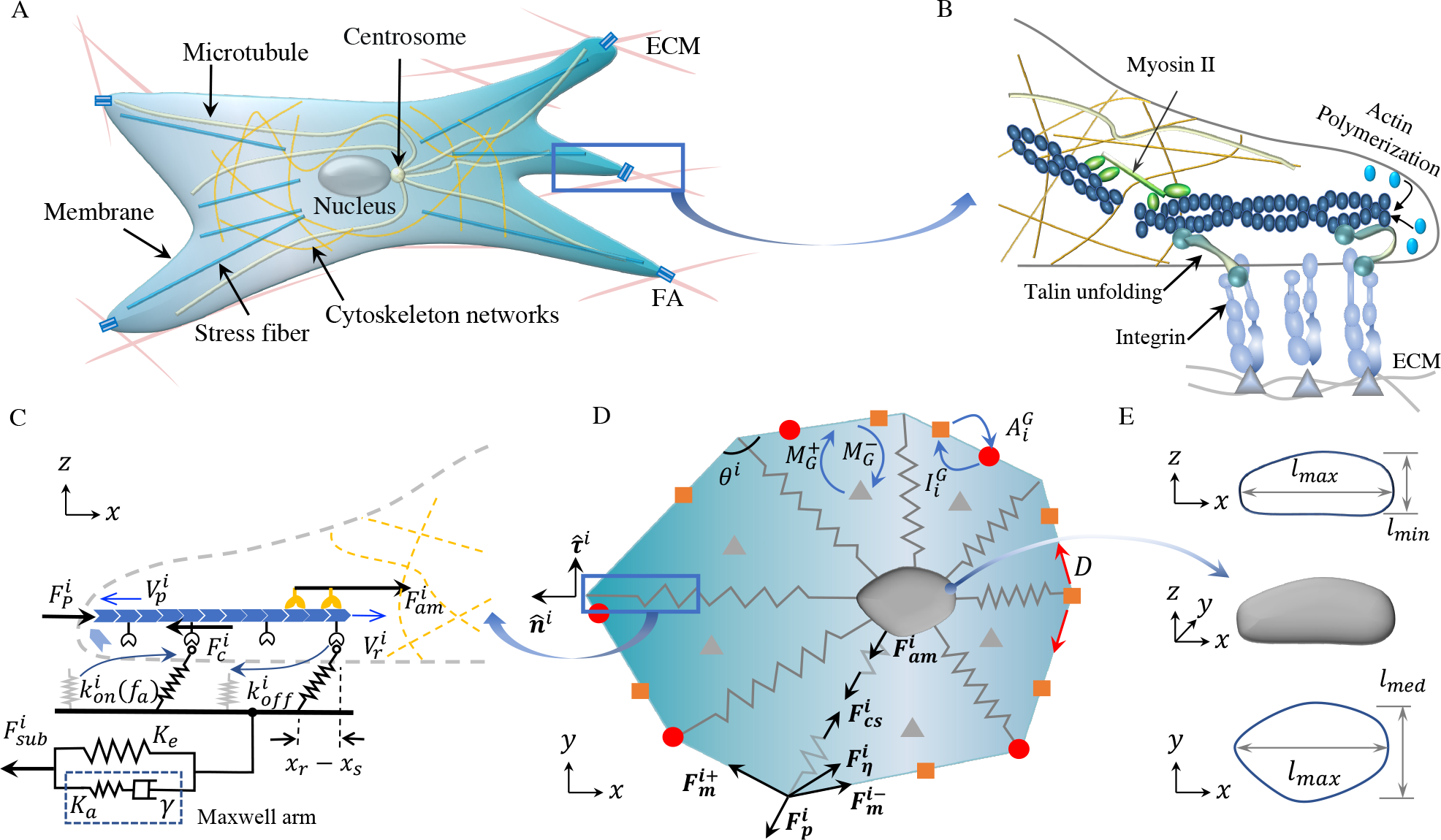
Multiscale whole-cell theory schematics. (A) Intracellular organization involved in mesenchymal migration. (B) Close-up view of the cytoskeleton–ECM linkage via integrins and the adaptor proteins (e.g., talin). (C) The parameters and variables of the motor-clutch model with adhesion reinforcement at each FA site. (D) The vertex-based model that couples the cell-matrix interactions at the subcellular scale to the chemomechanical dynamics at the cellular scale. The Rac1 concentration *R* and RhoA concentration *ρ* can each be in a membrane-bound active (red dots) or inactive state (orange squares), or dissolved in the cytoplasm (gray triangles). The conversion rates between these states are 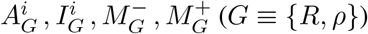, *D* is the diffusion convariable focal adhesion lifetimestant of the membrane-bound species (Eq. 15). The angle between the two membrane sections at a vertex is denoted by *θ*^*i*^. (E) The principal dimensions *l*_*max*_, *l*_*med*_, *l*_*min*_ of a deformed nucleus.

### 2.1 Equations of motion at subcellular scale

We first present the calculation of the cell vertex displacements due to the FA site dynamics and the mechanical balance at the corresponding vertex. For a vertex *i* with position vector **x**^*i*^(*t*), the local spreading velocity 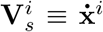 needs to be determined at every time step. In our model, the radial component 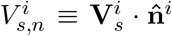 is controlled by the active processes, i.e., actin polymerization and actomyosin contraction regulated by the Rho GTPase proteins, while the polar spreading speed 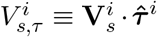 is set by the balance between the polar components of the passive forces due to the cell deformations (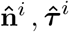 defined in Fig. 1 D).

The active radial spreading of a vertex with a speed 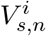 is fueled mainly by F-actin formation with a polymerization speed 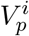 and counteracted by the retrograde G-actin flow with a speed 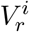, which yields

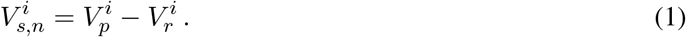

The polymerization rate at the vertex *i* is given by the ratio of the active Rac1 concentration 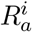 to its mean 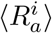 averaged over all vertices as 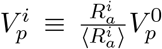, where 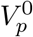 is the reference polymerization speed. The retrograde actin flow speed 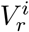 is promoted by the net resistance force against protrusion 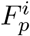 due to the cell membrane elasticity and myosin contractions (Fig. 1B, C) and impeded by an elastic restoring force 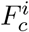 due to the formation of molecular bonds by proteins such as integrins, talin, and vinculin between F-actin and ECM. We thus propose a phenomenological relation

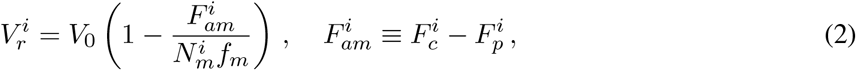

where *V*_0_ is the unloaded myosin motor speed, 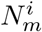 is the number of active myosin motors, and *f*_*m*_ is the force that stalls the activity of one myosin motor. Due to the increased iteration requirements for calculating 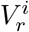 when reaching its minimum during loading-unloading focal adhesion cycles, we enforce the condition *V*_*r*_ = max(*V*_*r*_, 0) to ensure a non-negative retrograde velocity when convergence is not attained within a limited number of iterations (see Sec. S1 and Fig. S1). We have demonstrated that this condition does not affect the predictive accuracy of our whole-cell model (see Sec. S1 and Fig. S2). RhoA is known to induce myosin motor activation, leading to stress fiber formation and contractility [43, 44]. Therefore, we assume that the myosin motor number is controlled by the active RhoA concentration 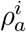 and its mean 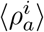 averaged over all vertices as 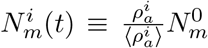, where 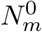 is the reference myosin motor number. Note that this assumption guarantees the constancy of the total motor number in the cell, in accordance with the conservation of mass [16, 26]. In Eq. 2, the elastic restoring force 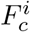 is determined from the FA dynamics with adhesion reinforcement, which is triggered by the stiffness of the viscoelastic substrate. On the other hand, the protrusion force 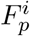 is set by the local force balance at each vertex in the presence of the nonlinear strain stiffening of the cytoskeleton. To calculate these two force strengths, we detail each of those processes next.

#### 2.1.1 Focal adhesion dynamics with adhesion reinforcement

To account for the adhesion reinforcement due to talin unfolding, we extend the augmented motor-clutch model for FA dynamics introduced in Refs. [21, 23] in order to calculate the clutch force 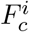 at each vertex. By denoting average displacements of all bounded clutches at the filament end by 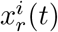 and the displacement of the substrate by 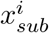, the engaged clutch is represented by a Hookean spring with tension 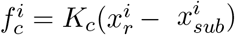 (*K*_*c*_ : spring stiffness) [45, 46]. At any instant *t*, the unbounded clutches must associate with a rate 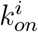. For the talin-low cells, a constant association rate 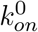 is typically assumed, corresponding to a constant clutch binding timescale 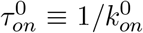 [17, 23, 28]. We introduce the adhesion reinforcement by assuming that, when the time-averaged clutch force 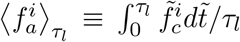 (*τ* : variable focal adhesion lifetime) is above a threshold force *f*_*cr*_, the clutch binding rate will increase per [21, 23],

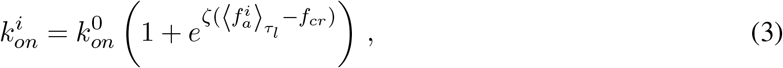

where *ζ* is a characteristic inverse force scale. Note that 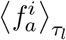 is determined by solving an isolated motor-clutch system separately (Eqs.2-6, see inset in Fig. S4 and Sec. S2), considering the updated protrusion force 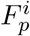 and the updated number of clutches 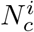 and motors 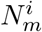 at vertex *i*. The clutch force 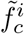 and the time 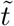 are variables in isolated motor-clutch modeling, distinct from the variables in our whole cell modeling. Due to the time dependence of 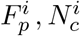, and 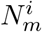, the variable 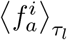 at every vertex must be updated at every *N* time steps. Our simulations have demonstrated that using *N* = 4000 time steps provides accurate predictions on cell migration speeds on stiff substrates, comparable to results obtained from more frequent updates of 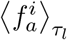 (see Fig. S5), while keeping computational costs reasonable.

Once formed, the molecular clutches must unbind at a dissociation rate 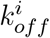 that depends on the clutch tension 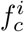. To that end, we use the functional form 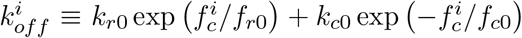. Here, 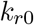 and 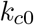 denote the unloaded off-rate and the unloaded catch-rate, respectively, *f*_*r*0_ is the characteristic rupture force, and *f*_*c*0_ is the characteristic catch force. It follows that the fraction of the engaged clutches (0 ≤ *P*^*i*^(*t*) ≤ 1) is governed by the mean-field rate equation [23, 45, 46],

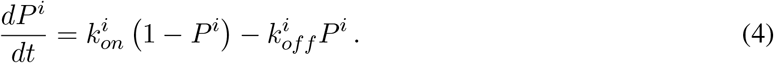

Denoting the number of available clutches as 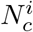, the total clutch force at the vertex *i* is then given by 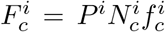. Here we incorporate the critical role of the Rac1 proteins in focal complex assembly by relating 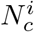 to the local Rac1 concentration, i.e., 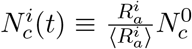, where 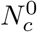 denotes the reference clutch number [47–49]. This assumption implies that the total count of clutches 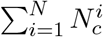 remains constant at any given moment, thereby adhering to the conservation of mass [16, 26]. The mechanical equilibrium condition at the cell-substrate interface demands that the total force sustained by the engaged clutches must be balanced by the substrate deformation, leading to

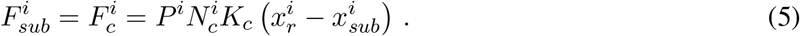

In Eq. 5, the substrate displacement 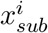 is an unknown to be determined from a constitutive model for the substrate viscoelasticity, which we focus on next.

#### 2.1.2 Constitutive model for substrate viscoelasticity

Our previous implementation utilized the classical Kelvin-Voigt model for the substrate relaxation dynamics [28]. However, since the Kelvin-Voigt model predicts a very rigid nonphysical behavior when *t < τ*_*r*_ (*τ*_*r*_ : substrate relaxation timescale) [50], it can result in a premature adhesion reinforcement. Comparatively, the standard linear solid (SLS) model allows for a better physical modeling by including a Maxwell arm (Fig. 1) [51]. It has also been demonstrated that the SLS model can capture the prominent relaxation timescale of the viscoelastic substrates fabricated by combining covalent and supramolecular crosslinking [23]. The SLS model expresses the constitutive relationship between 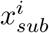 and the substrate force 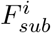 as (Fig. 1 C)

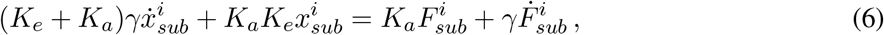

where *K*_*e*_ is the elastic stiffness at *t* → ∞, *γ* is the substrate viscosity, and *K*_*a*_ is the additional stiffness that governs the substrate relaxation with a timescale *τ*_*r*_ ≡ *γ/K*_*a*_. A stress relaxation test from a constant strain yields the instantaneous and long-term stiffness of the substrate as *K*_*t*→0_ = *K*_*e*_ + *K*_*a*_ and *K*_*t*→∞_ = *K*_*e*_, respectively. The instantaneous stiffness *K*_*t*→0_ characterizes the initial elastic response of the substrate when the viscous deformation and stress relaxation have not yet taken place in the limit *t* → 0. Thus, cells with a very short focal adhesion lifetime (*τ*_*l*_ *< τ*_*r*_) can only sense the instantaneous stiffness. In contrast, the long-term stiffness *K*_*t*→∞_ refers to the residual substrate stiffness after the viscous stresses have relaxed.

#### 2.1.3 Cytoskeletal stiffening

The passively deforming cell cytoskeleton, which consists of microtubules and intermediate filaments, is represented by multiple springs in our model (Fig. 1 D). Here we will assume that these cytoskeletal “springs” undergo strain-stiffening during large spreading events and thus exhibit nonlinear elasticity. This assumption is backed by the experiments performed, e.g., on NIH-3T3 fibroblasts that reveal a strong correlation between the cell rigidity and cell area during large spreading events [8]. Assuming this behavior is mechanically driven and thus must be generic across animal cells, we propose a phenomenological equation for the cytoskeletal stiffness *K*_*cs*_ in terms of the cell area *A*: Denoting the position vector of the cell nucleus by **x**_*nuc*_ and defining the length of a cytoskeletal spring as *r*^*i*^ ≡ |(**x**_*nuc*_ − **x**^*i*^)|, the cytoskeletal restoring force in our model is governed by the differential equation

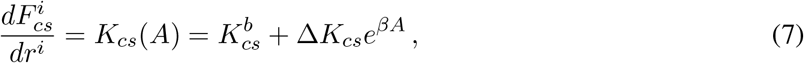

where 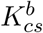 denotes the baseline stiffness. The term Δ*K*_*cs*_*e*^*βA*^ is introduced to describe the exponential hyperelastic behavior [52, 53], with the values of Δ*K*_*cs*_ and *β* obtained through linear regression of the experimental data in [8] (Sec. S3 and Fig. S6 A).

#### 2.1.4 Local mechanical equilibrium

We enforce local force balance at each vertex to determine the radial protrusion force 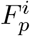 and the polar spreading speed 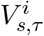. The cell membrane possesses a notable mechanical rigidity that enables it to endure a variety of stresses, which is vital for maintaining the integrity of cell structures [41, 54–56]. We model it as a closed loop of Hookean springs with stiffness *K*_*m*_ between neighboring vertices. The tensile forces acting on the *i*^*th*^ vertex can be expressed as 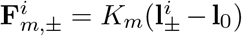, where 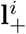 and 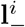 are the distance vectors between vertex *i* and its two neighbors, and **l**_0_ is the initial distance between two adjacent vertices. The drag force on vertex *i*, resulting from hydraulic resistance in the extracellular medium, can be expressed as 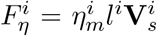, where 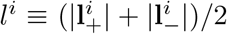 defines the average membrane length about the *i*^*th*^ vertex. The drag coefficient 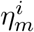 is proportional to the fluid viscosity but inversely proportional to the matrix permeability (see Sec. S4 for details) [57–60]. In addition to the membrane forces, each vertex experiences a protrusion force 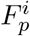 and a cytoskeletal radial force 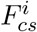. Thus, the net force balance at each vertex in the radial direction 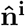 and the polar direction 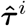 is given by

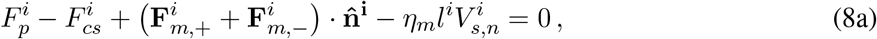

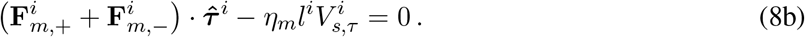

Eq. 8a and 8b yield 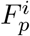 and 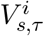, respectively. Altogether, Eq. 1–8b fully determine the vertex spreading velocities 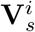 when the nucleus displacement **x**_*nuc*_ and the dynamical GTPase concentrations at each vertex are computed at the cellular scale.

### 2.2 Equations of motion at cellular scale

Next, we explain the global mechanical equilibrium that governs the nucleus motion and the intracellular Rho-GTPase dynamics, which directly influence the aforementioned subcellular processes.

#### 2.2.1 Viscous drag on deformed nucleus

Our model quantifies cell translocation by the net translation of the cell nucleus, which balances the forces between the cytoskeletal microtubules and intermediate filament bundles. At the cell periphery, these cytoskeletal complexes are linked to the F-actin at the FA sites, transmitting the net traction force from the extracellular matrix 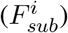 and the protrusion force from the cell boundary 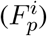 to the cell nucleus (Fig. 1). This tight linkage to the rest of the cell can deform the nucleus when the cell flattens under large spreading on stiff substrates [29]. To balance these peripheral and cytoskeletal forces, the nucleus undergoes viscous drag within the cytoplasm while it deforms under cell flattening. The viscous drag force on a particle is commonly described by the product of a drag coefficient, a characteristic particle size, and the velocity of the particle relative to the surrounding medium. Although the cytoplasm as the surrounding medium may undergo convective flows due to the intracellular biomolecular and organelle dynamics, its velocity around the nucleus can point in any direction at a certain time instant, either facilitating or opposing the movement of the nucleus at that instant. Thus, over extended periods of time, the contribution of the cytoplasmic flow to the motion of the nucleus can be neglected, and we treat the cytoplasm as a stationary viscous medium for the calculation of the nucleus velocity. In the case of a deformed nucleus with irregularities, we incorporate a shape factor denoted as *f*_*shape*_ into the analysis [61]. This leads to the following expression for the drag force:

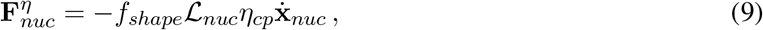

where *η*_*cp*_ represents the viscosity of the cytoplasm and ℒ_*nuc*_ is the characteristic particle size. For simplicity, we take 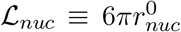 corresponding to Stokes’ flow as a first-order approximation to the cytoplasmic domain within finite cell height (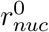: the initial radius of the nucleus). The shape factor *f*_*shape*_ quantifies the deviation of the nucleus from a perfect sphere with *f*_*shape*_ = 1. The Corey shape function can be used to determine *f*_*shape*_ based on the nucleus’s three principal lengths (Fig. 1 E) [61, 62],

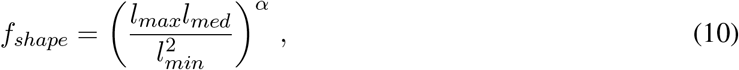

where the exponent *α* = 0.09 was obtained by fitting the experimental drag coefficient of non-spherical particles under Stokes flow [61]. For the deformed nucleus, the aspect ratio defined by the longest and the shortest dimensions can be related to the cell spreading area by 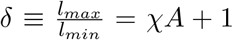, where the exponent *χ* = 0.0024 is obtained by fitting the experimental cell shape data in Ref. [63] (Fig. S6 B). The intermediate dimension *l*_*med*_ corresponds to the length of the minor axis of the nucleus on the *x*−*y* plane, and it was found to have a constant ratio to the length of the major axis in experiments (*l*_*med*_ = 0.8*l*_*max*_) [29]. Consequently, the nucleus shape factor can be simply related to the cell spreading area *A* by,

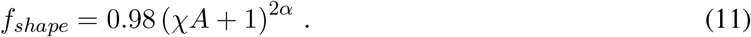

Experiments have demonstrated that the cytoplasm viscosity, like the strain stiffening, is strongly correlated with the cell spreading area [31, 32]. These experiments also indicate that the cell viscosity and cell stiffness exhibit the same trend with increasing substrate stiffness. Therefore, we assume an area-dependent cytoplasm viscosity (similar to that in Eq. 7) as

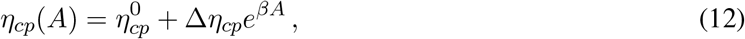

where 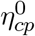 is the cytoplasm viscosity on soft substrates [64] and Δ*η*_*cp*_ controls the rate of viscosity increase with the cell spreading area *A*.

At the frame of the nucleus, the condition for the mechanical equilibrium between Eq. 9, the cytoskeletal and the subcellular forces

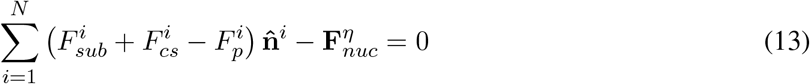

yields the nucleus migration velocity, 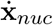, equivalent to the cell velocity in our model.

#### 2.2.2 Reaction-diffusion dynamics of GTPase concentrations

As with our previous work [28], here we adopt the biochemical reaction-diffusion equations introduced in [65, 66] to describe the dynamics of active and inactive GTPases. Since the active forms of the GTPases are predominantly associated with the cell membrane, which also serves as a major site for the conversion between active and inactive forms, we track the volume fractions of the signaling proteins in three forms [67, 68]: the active membrane-bound form 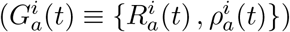, the inactive membrane-bound form 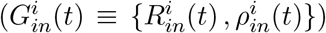, and the inactive cytosolic form 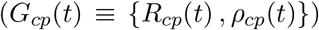. Given the rapid diffusion of inactive proteins in the cytosol, we assume *G*_*cp*_(*t*) remains uniformly distributed in the cytosol at all times. Both active and inactive membrane-bound forms diffuse with a diffusion constant *D* across the vertices. The corresponding diffusive fluxes are given by the Fick’s law in a finite difference formulation as

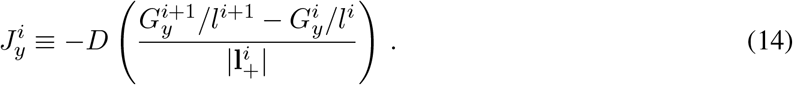

Eq. 14 takes into account the effect of the deformed cell shape on the diffusive flux by updating the vertex coordinates at each time step. Different forms of the proteins on a vertex are interconvertible with the inactive-to-active rates 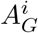, active-to-inactive rates 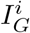, and inactive-to-cytosolic association and disassociation rates 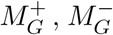 (Fig.1 D). Altogether, the reaction-diffusion kinetics is governed by

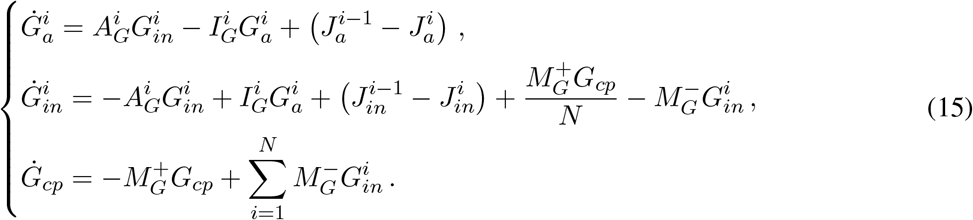

The total amounts of Rac1 and RhoA are constant due to the conservation law 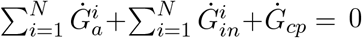. The active Rac1 and RhoA GTPase volume fractions 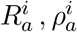 regulate the cell migration by controlling the actin polymerization speed 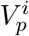 (Eq. 1), the number of myosin motors 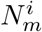 (Eq. 2), and the clutch number 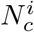 at each FA site (Eq. 5). Different from Refs. [65, 66], the rate terms 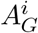 in our formulation account for the reverse coupling from the cell deformation to the signaling pathways in addition to the mutual inhibition of the Rac1 and RhoA (Sec. A1). Since the filopodial protrusions of a mesenchymal cell grow and shrink at timescales comparable to the migration times, this chemo-mechanical feedback prevents the Rac1 and RhoA dynamics from reaching a steady bistable polarized state. Instead, when the transient chemical polarity ceases, our simulation algorithm reinstates the polarization stochastically to sustain the random migration patterns in many mesenchymal phenotypes [26, 27].

### 2.3 Simulation procedure

We represent the initial vertex configuration of the cell as a regular 16-sided polygon due to its decent convergence properties [28]. All initial forces, velocities, displacements, and substrate deformations are set to zero. Since chemical signaling drives cell polarization followed by directional locomotion on uniform substrates [28], a nonuniform initial distribution of the active Rac1 protein volume fraction 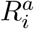 is enforced with a higher value at the cell front. Likewise, a polarized initial distribution of the active RhoA protein volume fractions 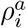 is taken with accumulation at the cell rear (manual polar symmetry breaking). In contrast, the initial conditions for 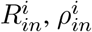 are uniform at time *t* = 0, and the cytoplasmic form *R*_*cp*_, *ρ*_*cp*_ can be determined from the total volume fraction conservation (Fig. S9). For directed migration on nonuniform substrates, uniform initial signaling distributions must be assumed since the polarity will be set by the gradients of the ECM stiffness or viscosity. Starting from these initial conditions, the vertex spreading and the nucleus velocities are calculated by solving Eq. 1–13, and the Rho-GTPase volume fractions are updated by solving Eq.14, 15, A1, A2 by following the algorithm in Fig. S4. When the cytoplasmic inactive Rac1 volume fraction *R*_*cp*_ reaches a steady state, we reinitialize 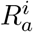 and 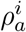 stochastically to mimic the random nature of the protrusion formations in the mesenchymal cells. To incorporate the adhesion reinforcement (Eq. 3), we fix *ζ* = 0.5*pN* ^−1^ since it leads to a good agreement between the simulated spreading areas and the experimental data in Ref. [8] (Fig. S10). The parameters Δ*K*_*cs*_ = 2.8 *pN/μm* and *β* = 0.00185 (Eq. 7) and *χ* = 2.4 × 10^−3^ (Eq. 11) provide the best fit to the experimental data from Ref. [8] and are kept constant in all simulations (Fig. S6). All fixed simulation parameters are listed in Table S1.

We present the simulation results in dimensionless form by introducing a position scale *L* ≡ 2*πr*_0_ = 10*π μm*, speed scale *V*_0_ = 120 *nm/s*, migration timescale *T* ≡ *L/V* ≈ 260 *s*, and force scale 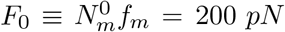. For each simulation, we run *M >* 10^6^ time steps with a time step of approximately Δ*t* = 1.5 × 10^−3^ *s* (Δ*t* ≈ 5.7 × 10^−6^ in dimensionless units), corresponding to the migration dynamics over at least an hour in real units. The spreading area *A* is calculated as the average cell area over the last 10^6^ time steps in a simulation. We define the migration speed *V* as the nucleus trajectory length *X* ≡ |**x**_*nuc*_(*M* Δ*t*) − **x**_*nuc*_(0)| divided by the total time *t*_*total*_ ≡ *M* Δ*t*, i.e., *V* ≡ *X/t*_*total*_. The ranges of the ECM material parameters used in the simulations are listed in Table 1. For each set of the ECM parameters, we run *n* simulations to compute the sample mean spreading area, Ā, and the mean migration speed, 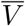. The effects of the ECM viscoelasticity on the spreading area, migration speed, and migration direction are validated by the experimental data taken from Refs. [6, 16, 27].

**Table 1:** Residual substrate stiffness *K*_*e*_, additional stiffness *K*_*a*_, and viscosity *γ* ranges in simulations.

| Parameter | Real units | Dimensionless units |
| --- | --- | --- |
| $K_e$ | 0.1-100 $pN/nm$ [17, 23, 46] | $5\pi - 5000\pi$ |
| $K_a$ | 0.1 – 20 $pN/nm$ | $20\pi - 4000\pi$ |
| $\gamma$ | 0.01 – 100.0 $pN \cdot s/nm$ [23] | 0.06 – 60 |

## 3 Results

### 3.1 Simulations reproduce experimental cell spreading and migration speed dependence on substrate stiffness

With increasing stiffness *K*_*e*_ on an elastic substrate (*γ* = 0, *K*_*a*_ = 0), the cell spreading area increases monotonically, in agreement with the experiments on the adhesion-reinforced U373-MG human glioma cell spreading on polyacrylamide hydrogels (Fig. 2 A) [6]. This can be understood by inspecting the FA dynamics at the subcellular scale: According to Eq. 2, 5, the cell spreading speed at a vertex 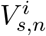 is set by the competition between the polymerization speed 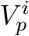 and the actin retrograde flow speed 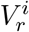, which is counteracted by the stiffness-dependent focal traction force (Sec. A2)

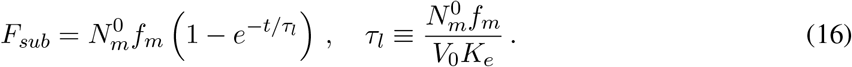

**Figure 2:**
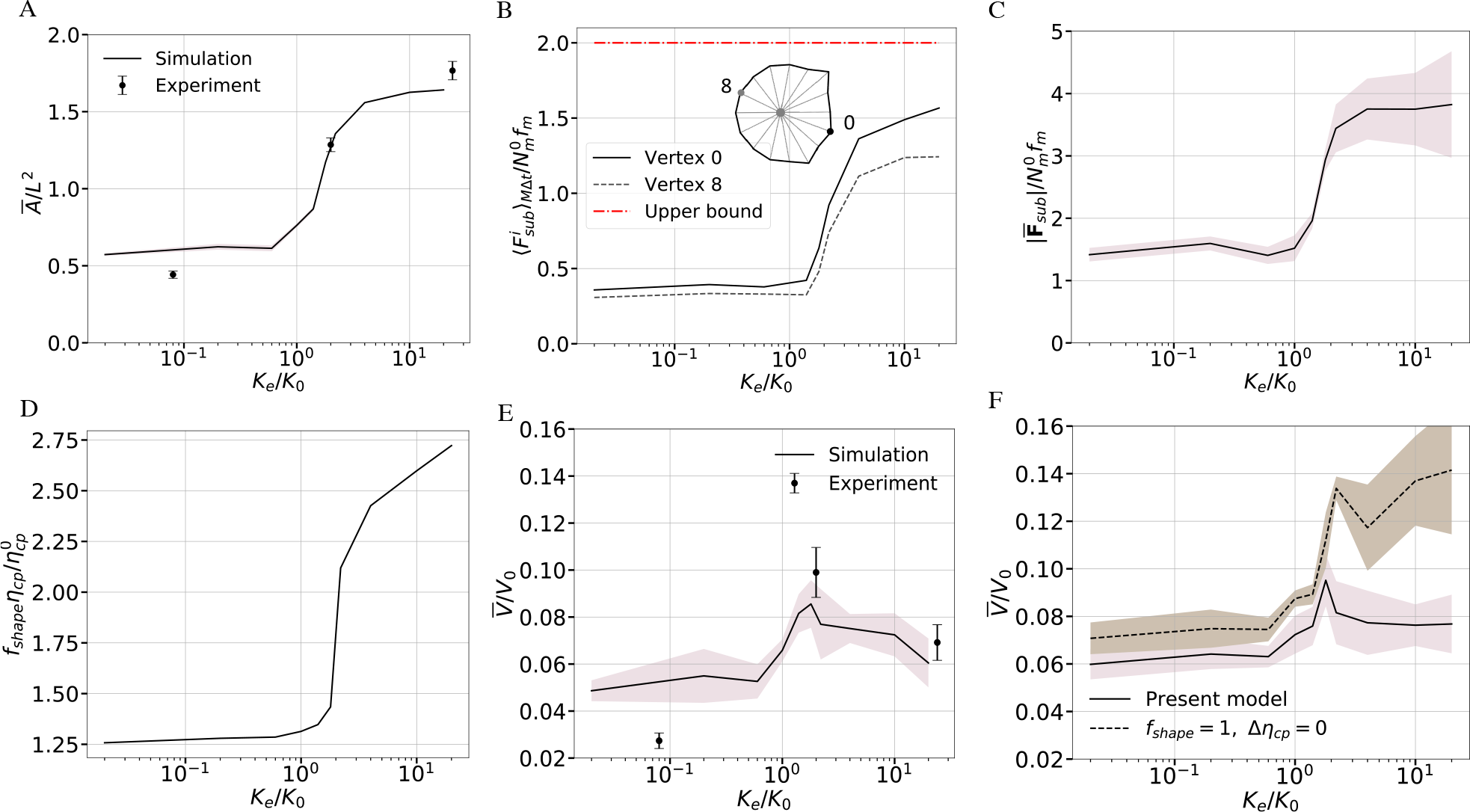
Cell spreading area and migration speed on elastic substrates. (A) The dimensionless mean cell area Ā */L*^2^ (full line) as a function of the dimensionless substrate elastic stiffness *K*_*e*_*/K*_0_. For comparison, the dots and error bars are the experimental data from Ref. [6] [69]. (B) The average dimensionless substrate traction force versus *K*_*e*_*/K*_0_ at vertex 0 (cell front) and vertex 8 (cell rear). The red dash-doted line gives the upper bound of the substrate traction force on very stiff substrates. The calculation of the averaged substrate traction forces 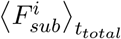 from the full focal adhesion cycles is illustrated in Fig. S11. (C) The dimensionless mean and standard deviations of the net traction force versus *K*_*e*_*/K*_0_. The calculation of the time-averaged net traction force is illustrated in Fig. S12. (D) Cell-spreading induced nucleus viscous drag increase 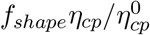 versus *K*_*e*_*/K*_0_, which is calculated by using Eqs. 9, 11, 12 for the mean spreading area values in A. (E) The simulated dimensionless mean migration speed 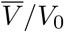 versus *K*_*e*_*/K*_0_ (full curve) and the experimental data for the U373-MG human glioma cells (dots and error bars) [6]. (F) The influence of the shape factor *f*_*shape*_ of the deformed nucleus and the area-dependent cytoplasm viscosity *η*_*cp*_ on the migration speed (Eq. 11, 12). The mean values or the standard deviations are calculated over *n* = 10 simulations at each data point in A, C, D, E, F.

Eq. 16 is derived for an isolated motor clutch system where the engaged clutch fraction is assumed to be saturated on a compliant substrate 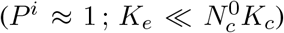. For sufficiently low *K*_*e*_ where the adhesion reinforcement mechanism is not triggered (see Eq. 3), the traction force starts building up at an FA site after the molecular clutches form at a timescale 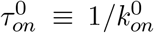, and develops with a characteristic time *τ*_*l*_ (Eq. 16). The comparison between the two timescales 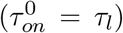 defines a threshold stiffness 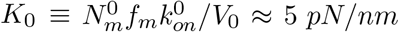 (Table 2). On soft substrates (*K*_*e*_ ≤ *K*_0_), since the force build-up time *τ*_*l*_ is much longer than the clutch formation time *τ*_*on*_, the adhesion force mostly remains low, i..e, 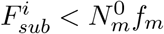 (Eq. 16; Fig. 2 B). For a stiff substrate with *K*_*e*_ *> K*_0_, the decrease in the force build-up time *τ*_*l*_ must cause a steep increase in the average force per bond 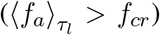, triggering the adhesion reinforcement mechanism with an augmented binding rate *k*_*on*_ (Eq. 3, Fig. S3). The reinforced binding rate can even saturate the bounded clutches on a very stiff substrate, i.e., *P*^*i*^ → 1 (Fig. S3 F). Moreover, a higher 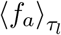 on stiff substrates lead to higher disassociation rates *k*_*off*_, which must balance the binding rate in Eq. 4 to reach a steady bound clutch fraction *P*^*i*^ ∼ 1. This condition yields an estimate for the instantaneous force per bond as 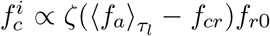 (Sec. A3). This allows us to estimate an upper bound of the substrate traction force on a stiff substrate as 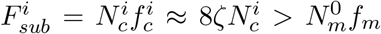, in agreement with the simulations (Fig. 2 B). This elevated traction at *K*_*e*_ *> K*_0_ across the cell is the main driving factor behind enhanced cell spreading. Furthermore, the Rho GTPase polarization leads to a higher traction force at the cell front than at the rear since the motor clutch number 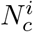 is controlled by the Rac1 concentration (Fig. 2 B). Due to the chemically induced polarization, the cell acquires a larger net traction force on stiff substrates than on soft ones (Fig.2 C).

**Table 2:**
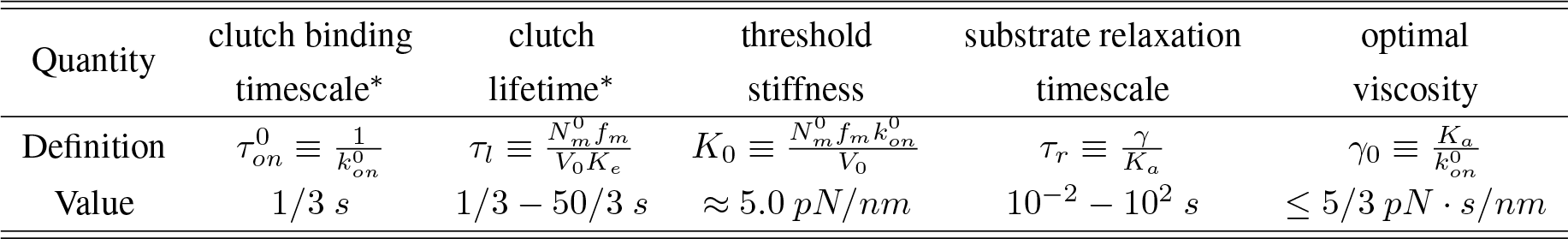
Characteristic scales for spreading and migration dynamics. For a viscoelastic substrate, *K*_*a*_ is the additional stiffness, and *γ* is the viscosity in the SLS model. Additionally, 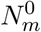 is the reference myosin motor number, *V*_0_ is the unloaded myosin motor speed, *f*_*m*_ is the force that stalls the activity of one myosin motor, and 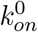 denotes the reference clutch binding rate. The scales denoted by ^*^ are valid for substrates with a stiffness lower than *K*_0_.

The cell motility is governed by the competition between the net traction force and the viscous drag of the nucleus since the remaining forces are negligible in Eq. 13 (Fig. S13). Because large cell spreading on stiff substrates can deform the nucleus (Fig. 2 D) and alter the cytoplasmic viscosity (Eq. 12), the viscous drag force on the nucleus must increase, resisting the net traction force. Consequently, the migration efficiency decreases beyond the threshold stiffness *K*_0_, creating a non-monotonic stiffness-speed profile, which also quantitatively reproduces the U373-MG human glioma cell speed on polyacrylamide hydrogels (Fig. 2 E) [6]. To consider the effect of potential cytoplasmic flows to the nucleus motion, we made a first-order approximation by linking the drag force to the relative velocity between the nucleus and the cytoplasm. Fig. S8 demonstrates that this approximation do not substantially influence the efficiency of cell movement over long times, supporting the validity of Eq. 9. In our simulations, ignoring the nucleus shape change and assuming constant cytoplasmic viscosity recovers a monotonic relationship between the migration speed and substrate stiffness, which validates their role in the non-monotonic speed-stiffness relation (Fig. 2 F). Furthermore, we have performed parametric studies to explore the relative impact of the nucleus shape change and the cytoplasmic viscosity increase on the non-monotonic speed-rigidity relationship (Fig. S14). We found that while both factors influence the migration speed on soft substrates, the effect of cytoplasmic viscosity increase becomes more significant on stiff substrates, emphasizing its critical role in capturing the non-monotonic relationship.

### 3.2 Substrate viscosity affects cell spreading area and migration speed

We next discuss the effect of substrate viscosity on the cell area and migration speed on a substrate that is soft in the long time limit, *K*_*e*_ = 0.1 *pN/nm*. When *K*_*e*_ + *K*_*a*_ *< K*_0_ (soft regime), the adhesion reinforcement effect can be neglected even on viscous substrates. However, when *K*_*e*_ + *K*_*a*_ ≥ *K*_0_ (stiff regime), a large viscosity may trigger the adhesion reinforcement regime and alter the spreading behavior. We discuss both regimes below.

On substrates with a low instantaneous stiffness (*K*_*e*_ + *K*_*a*_ *< K*_0_), the cell spreading area varies in a non-monotonic manner with increasing viscosity (Fig. 3 A, B). This can be explained by examining the FA dynamics at the subcellular scale, similar to the cell spreading on elastic substrates. At an FA site without adhesion reinforcement, time averaging the traction force (see Fig. S15 A, B) over a clutch lifetime *τ*_*l*_ leads to (Sec. A2)

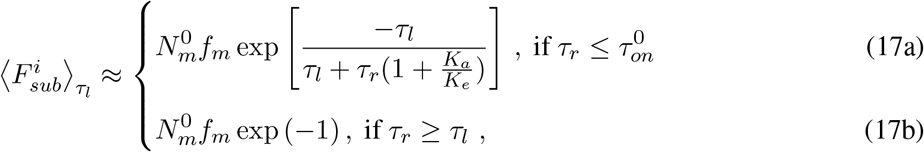

where the timescales *τ*_*l*_, 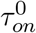, *τ*_*r*_ are defined in Table.2. Eq. 17a indicates that the average traction force at an FA site must increase with increasing viscosity when 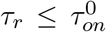. At 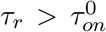, since slower substrate relaxation leads to the FA lifetime reduction when (Fig. S15 C), the average traction force should exhibit a decreasing profile with a lower bound that is estimated in Eq.17b. That way, Eq. 17 indicate the presence of a non-monotonic traction force-viscosity profile in an isolated motor-clutch system on soft substrates. Note that when 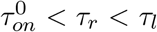, the average substrate force is not amenable to an analytical approximation. Still, given the biphasic profile suggested by Eq. 17, it is plausible to assume that the average substrate force reaches a maximum when 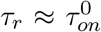, equivalently at an optimal viscosity 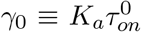 (Fig. S15 C), thereby maximizing the cell spreading area. To further validate our model, we simulated spreading areas of human fibrosarcoma cells HT-1080 on viscoelastic substrates consisting of interpenetrating alginate networks (IPN) [27]. The experimental material parameters of the substrates with different relaxation rates were obtained by data fitting to the stress relaxation tests in Ref. [27] (Sec. S3 and Fig. S16). Clearly, our whole-cell model accurately replicates the experimentally observed spreading areas of the HT-1080 cells on soft substrates with different stress relaxation rates (Fig. 3 C), indicating that cell spreading is suppressed with increasing viscosity when 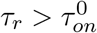.

**Figure 3:**
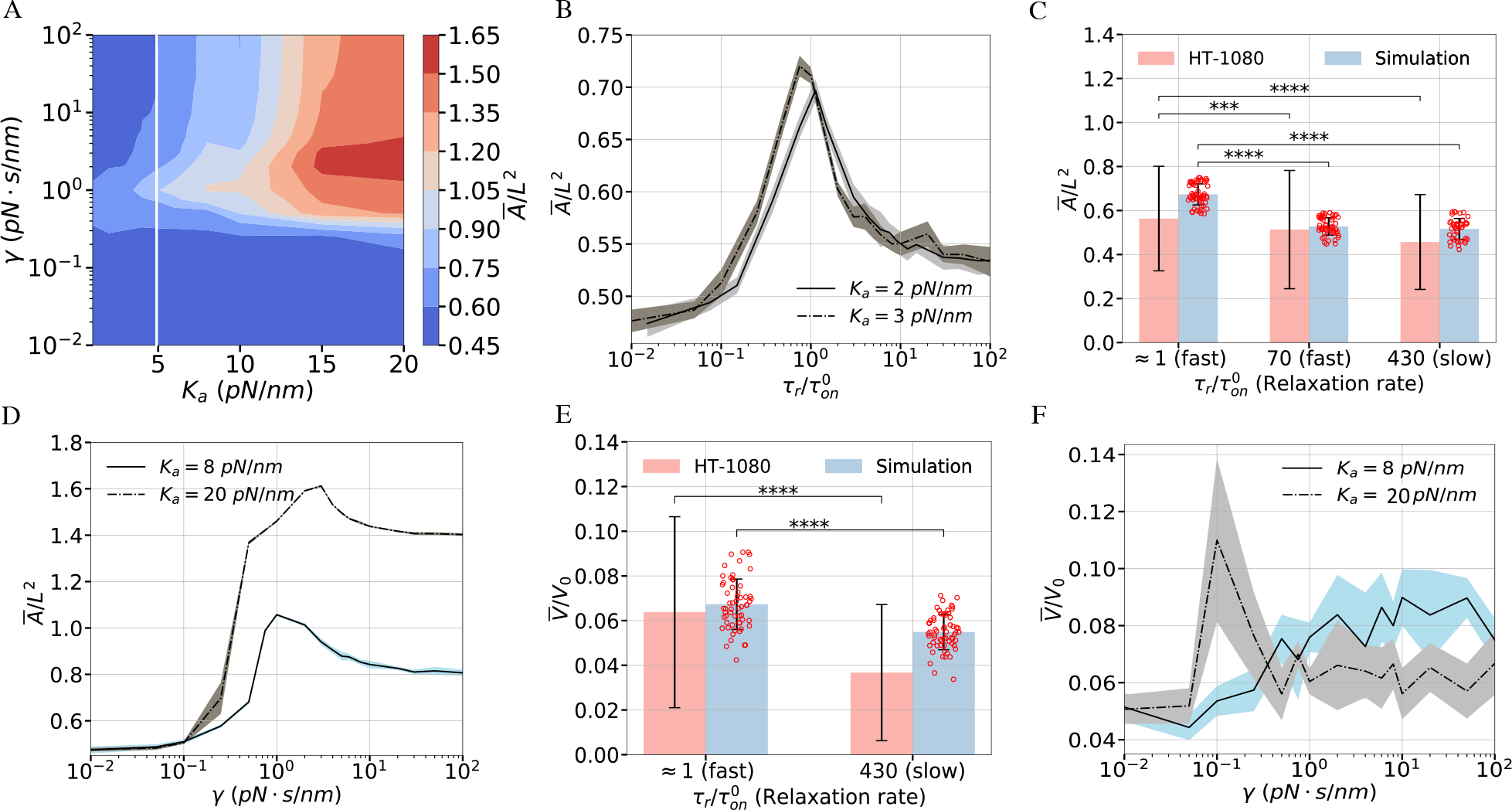
Effect of substrate viscosity on spreading area and migration speed. (A) Contour plot of the mean dimensionless cell area Ā*/L*^2^ on viscoelastic substrates as a function of the additional stiffness *K*_*a*_ and substrate viscosity *γ* (*K*_*e*_ = 0.1 *pN/nm*). The white line denotes the threshold stiffness *K*_0_. (B) Ā*/L*^2^ and its standard deviation (shaded area) versus the ratio of the material relaxation timescale *τ*_*r*_ ≡ *γ/K*_*a*_ to the clutch binding timescale 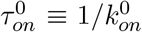 on soft substrates. (C) For a fixed instantaneous stiffness *K*_*e*_ + *K*_*a*_ = 2.6 *pN/nm*, the simulated spreading area Ā and its standard deviation (*n >* 50) and experimental data for HT-1080 cells (data from [27]). By fitting the stress-time data in the stress relaxation test, the viscosities of the fast, intermediate (med), and slow relaxing substrates are found as *γ*_*fast*_ = 1 *pN s/nm, γ*_*med*_ = 60 *pN s/nm, γ*_*slow*_ = 360 *pN s/nm* (Fig. S16). (D) The dimensionless spreading area Ā*/L*^2^ and its standard deviation as a function of *γ* for different *K*_*a*_ *> K*_0_. Since the clutch binding timescale *τ*_*on*_ depends on the adhesion reinforcement on stiff substrates, Ā*/L*^2^ is plotted against the substrate viscosity *γ*. (E) The mean migration speed 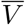 and its standard deviation in the simulations (*n >* 50) versus experiments of HT-1080 cells (data from [27]) on fast-relaxing and slow-relaxing soft substrates, for which the material parameters are extracted from fitting the stress-time data (Fig. S16). (F) The dimensionless mean migration speed 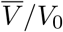 and its standard deviation as a function of *γ* for two additional stiffness values *K*_*a*_ = 8 *pN/nm* and *K*_*a*_ = 20 *pN/nm*. The contour plot in A contains 180 data points. The mean and standard deviations are calculated with *n* ≥ 5 simulations at each point in B, D, F. In C and E, statistical analyses were conducted using the t-test to compare simulation results with experimental data. Levels of statistical significance (*p*) are indicated as follows: * * ** : *p <* 0.0001, and * * * : *p <* 0.001.

At a larger instantaneous substrate stiffness *K*_*e*_ + *K*_*a*_ ≥ *K*_0_, the cells only perceive the long-term stiffness *K*_*e*_ and maintain small spreading areas at a low substrate viscosity *γ* (Fig. 3 D). For higher *γ*, the slowly relaxing substrate maintains the stiffness *K*_*e*_ + *K*_*a*_ ≥ *K*_0_ at later times, sustaining the adhesion reinforcement regime and thus promoting cell spreading. Moreover, because the adhesion reinforcement effect becomes more pronounced on stiffer substrates, a larger additional stiffness, such as *K*_*a*_ = 20 *pN/nm* further increases the cell spreading area at high *γ* (Fig. 3 D).

As opposed to the spreading area that can be understood at the single FA level, the change in the cell migration speed as a function of *γ* must be examined by virtue of the net traction force 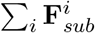 at the cellular scale. At an instantaneous substrate stiffness *K*_*a*_ + *K*_*e*_ *< K*_0_, where the adhesion reinforcement is not activated, the high viscosity of *γ > γ*_0_ can result in a decrease of the FA lifetime and thus a lower traction force (Sec. A2 and Fig. S15 D). Our simulations reveal that migration on fast relaxing substrates 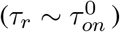 is more efficient than on slow relaxing substrates 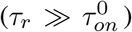 (Fig.3 E). This is in quantitative agreement with the migration speeds of the MDA-MB-231 human breast cancer adenocarcinoma cells on fast and slow relaxing IPN [27]. A more comprehensive parametric study demonstrates the presence of a biphasic speed-viscosity relation (Fig. S17 A, B). The optimal viscosity corresponding to the maximum migration speed tends to be slightly higher than *γ*_0_ (for maximum cell spreading), primarily due to the prolonged FA lifetime at the front of the cell (Sec. A2). On substrates with high instantaneous stiffness (*K*_*a*_ + *K*_*e*_ ≥ *K*_0_), the net traction force increases with viscosity due to the adhesion reinforcement effect (Fig. S17 C). Since cells have limited spreading areas due to the weak engagement of the adhesion reinforcement regime at *K*_*a*_ = 8 *pN/nm* (Fig. 3 D), the change in the nucleus drag force is marginal. Thus, we obtain a nearly monotonic increase in the migration speed with viscosity (Fig. 3 F). However, with a very large additional stiffness of *K*_*a*_ ≫ *K*_0_, strong adhesion reinforcement on the viscous substrate results in very large spreading areas (Fig. 3 D), which significantly increases the nucleus drag forces and in turn leads to a biphasic speed-viscosity profile (Fig. 3 F).

### 3.3 Migration direction is set by stiffness or viscosity variation on nonuniform substrates

Having elucidated the effect of substrate stiffness and viscosity on cell area and migration speed, we investigate the sensitivity of cell migration to a stiffness gradient on elastic substrates or a viscosity gradient on viscoelastic substrates. To eliminate the effect of chemical polarity on migration direction in the simulations, we specify the initial conditions of a chemical apolar cell by assigning a random distribution of the active Rac1 concentration 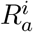 and setting the other membrane-bound Rho GTPase concentrations 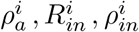 constant (Fig. S18). Migrating cells exhibit limited migration speeds 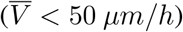, leading to a substantial amount of time required for simulating cell locomotion over long distances. Therefore, for the sake of computational efficiency, we adopt a 200*μm* × 300*μm* domain at about a 1 : 10 scale of the experimental platform Fig. 4 A). We compared our migration simulations of 83 individual cells on elastic gradient substrates to the experimental migration patterns of the MDA-MB-231 cells on polyacrylamide hydrogels with a stiffness gradient 0.5 − 22.0 *kPa* [16]. We record the migration trajectory of each simulated cell for 7.2 hours, which is 1*/*10 of the experimental observation time (72 *h*). Due to the difficulty in quantifying the substrate gradient in the experiments, we assume a constant gradient ∇*K*_*e*_ ≈ 72 *pN/μm*^2^ in the +*y*−direction in the simulations; non-dimensionalization by the initial cell diameter 2*r*_0_ and the threshold stiffness *K*_0_ yields the unitless gradient 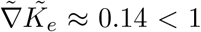. The simulated cell trajectories indicate that all cells with random locations at *t* = 0 move up the stiffness gradient regardless of their initial positions, i.e., the cells undergo robust durotaxis (Fig. 4 A). Also, the percentages of the simulated cells in three substrate portions demonstrate the strong migration trend towards the stiffest region (Fig. 4 B), in good agreement with the gradient sensitivity of the MDA-MB-231 cells with talin- and vinculin-mediated FA formations [16]. We further confirm that almost all cells eventually migrate to the stiffest region after *>* 12 *h* (Fig. 4 B). The robust durotaxis behavior is mainly due to the monotonic increase of the traction force as a function of the substrate stiffness at an FA site due to the adhesion reinforcement (Fig.2 B). In other words, the FA sites attached to the stiffer regions generate a higher traction force than those attached to the softer regions on a nonuniform substrate, driving the cell up the stiffness gradient.

**Figure 4:**
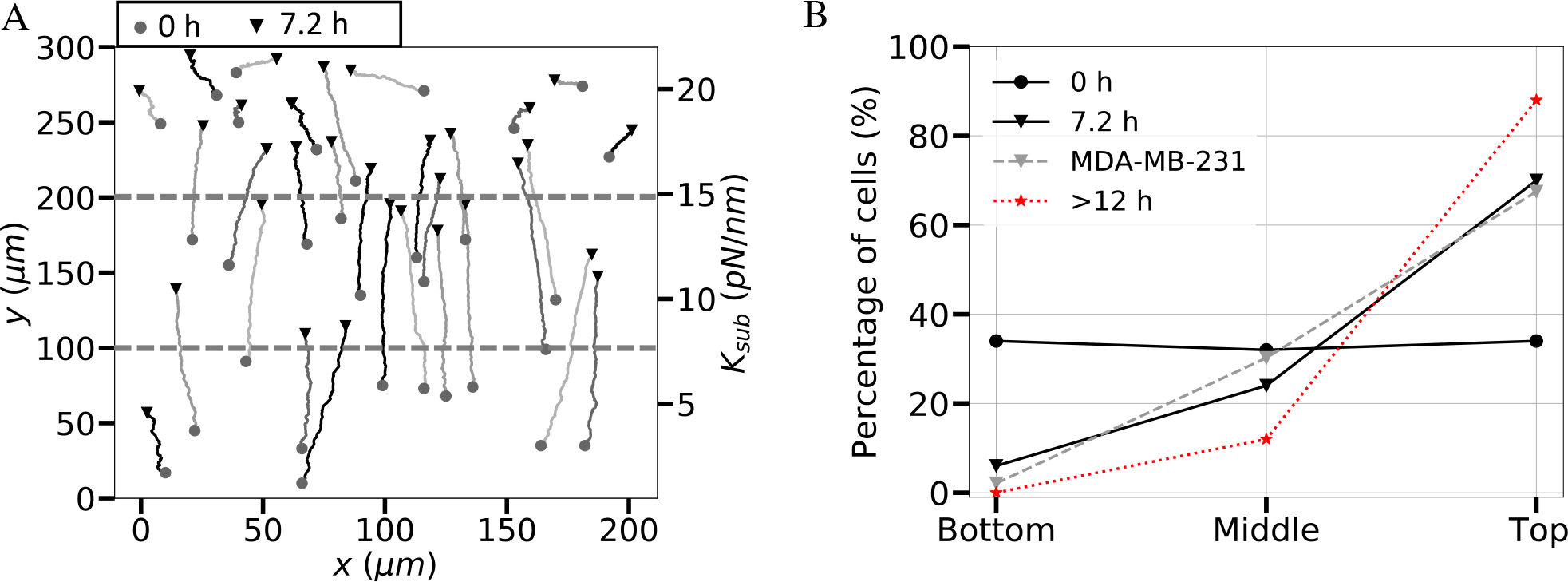
Simulations reproduce cell durotaxis on elastic gradient substrates. (A) Trajectories of 30 simulated cells over *>* 7.2 hours in real units on an elastic substrate with a linear stiffness variation between *K*_*e*_ = 0.5 − 22 *pN/nm* in the *y* − direction. (B) The percentage of the simulated cells out of *n* = 83 simulations on the bottom-, middle-, and top-third of the gradient substrate as a function of time, i.e., at *t* = 0 *h* (initial condition), *t* = 7.2 *h, t >* 12 *h*, which are compared to the distributions of the MDA-MB-231 cells (gray dashed lines) that are seeded on polyacrylamide hydrogels with a stiffness gradient and observed for 72 hours [16].

We next sought to understand the impact of substrate stress relaxation on the directed migration of cells on soft substrates. In our simulations, the two stiffness values in the SLS model are set to be *K*_*a*_ = 2.9 *pN/nm* and *K*_*e*_ = 0.1 *pN/nm*. We assume a nonuniform viscosity *γ < γ*_0_ ≈ 1 *pN* • *s/nm* with a constant gradient ∇*γ* = 12.5 *pN* · *s/μm*^2^ in +*x*−direction, equivalent to 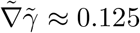 in the unitless form (non-dimensionalized by *γ*_0_ and 2*r*_0_). For computational efficiency, we used a 80 *μm* × 40 *μm* domain, which still much larger than a migrating cell size in ±*x*−direction.The simulations demonstrate that cells migrate from the soft elastic region (*γ* → 0) to the viscoelastic region, which is also evidenced by the biased angular displacement towards the +*x*−direction (Fig. 5 A, B). The same trend was observed in human mesenchymal stem cells that migrate towards regions with a larger relaxation modulus on a collagen-coated polyacrylamide gel. This behavior is referred to as viscotaxis [39]. Once the substrate viscosity surpasses the threshold (*γ > γ*_0_), cells migrate against the viscosity gradient with the angular displacements biased toward faster relaxation regions, which can be referred to as “anti-viscotaxis” (Fig.5 C, D). As a control, the random cell trajectories and unbiased angular displacements were observed on a uniform viscoelastic substrate (Fig. 5 E, F). These findings suggest that a viscosity gradient draws the cells to a region with 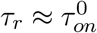. The underlying mechanism for the dependence of the migration direction on the viscosity gradient can be explained by inspecting the biphasic viscosity - traction force relationship at an FA site. On soft substrates with a relaxation timescale 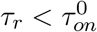, an increasing viscosity contributes to an enhanced substrate traction force that counteracts retrograde actin flow. This leads to a substrate-induced polarity along the gradient. In contrast, slower relaxing substrates 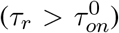 lead to decreasing clutch lifetime (Fig. S15 C) and the reduced mean traction forces (Eq.17). As a result, the edges of the cell attached to the faster relaxing substrate regions gain more traction, navigating the cell against the viscosity gradient (Fig. 5 G).

**Figure 5:**
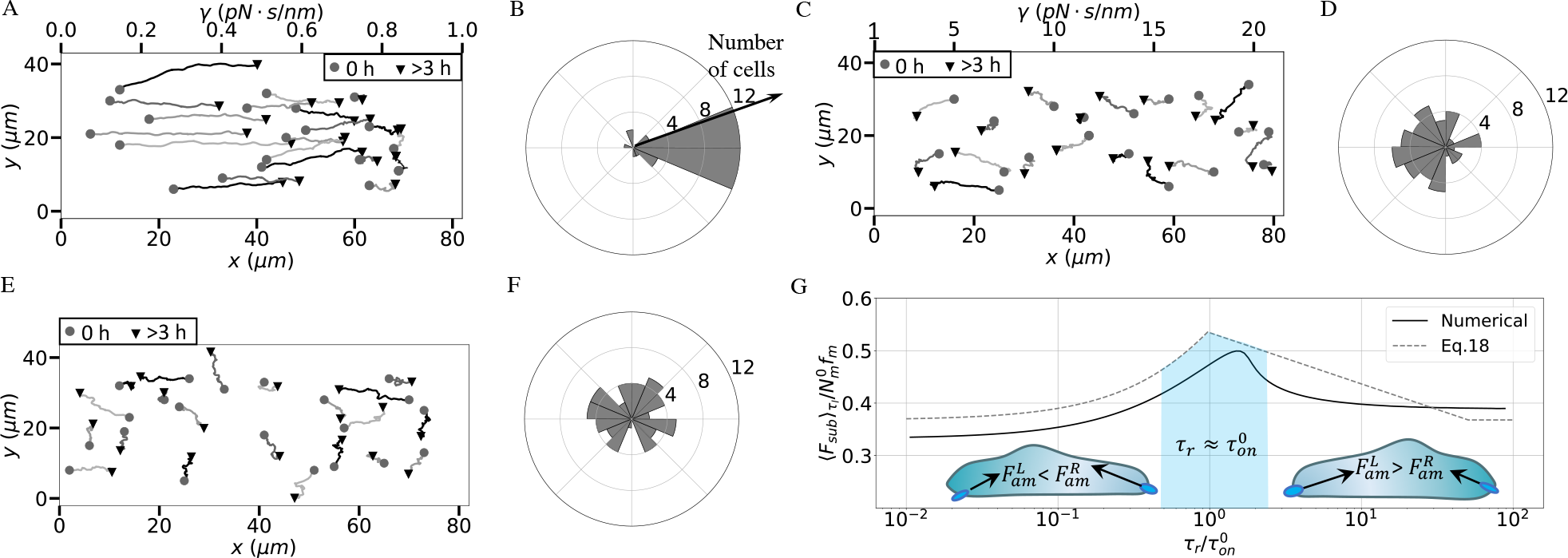
Directed migration is regulated by viscosity gradient on soft substrates. Representative trajectories of 20 cells and angular displacements of ≈ 50 cells on a viscoelastic substrate (*K*_*a*_ = 0.1 *pN/nm, K*_*e*_ = 2.9 *pN/nm*) with (A), (B) a linearly varying viscosity *γ* = 0.0 − 1 *pN* • *s/nm*, (C), (D) *γ* = 1.0 − 20 *pN* • *s/nm*, and (E), (F) a uniform viscosity *γ* = 10*pN* • *s/nm*. (G) The simulated versus analytical traction forces (averaged over *τ*_*l*_) as a function of the ratio of the material relaxation timescale *τ*_*r*_ ≡ *γ/K*_*a*_ to the clutch binding timescale 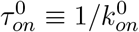 (Table 2). The traction forces at the two opposite ends of a cell along the direction of the motion are labeled by 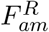 or 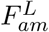, respectively.

## 4 Discussion

Our multiscale theory involves a detailed description of the adhesion reinforcement regime at each FA site and includes the cytoskeleton strain-stiffening effect, which quantitatively reproduces the large cell spreading areas on stiff substrates. Importantly, by incorporating the viscous drag force increase due to the nucleus shape change and nonlinear cytoplasmic viscosity as a function of the spreading area, we have reproduced the non-monotonic dependence of the migration speed on substrate stiffness. Our findings indicate that the interplay among larger spreading areas on stiff substrates, variable cytoplasmic viscosity, and nucleus flattening can influence the drag forces in cell migration and give rise to the non-monotonic speed profile. Therefore, the mechanism underlying the non-monotonic speed-stiffness relation in this study is distinct from the biphasic speed-stiffness relation discussed in our previous work [28]. The biphasic speed profile in [28] is attributed to the biphasic dependence of the focal adhesion force on substrate stiffness in cells that lack an adhesion reinforcement. A limitation of our previous model was that it cannot reproduce the monotonic increase of spreading area with stiffness. In contrast, our current model effectively reconciles both the observed monotonic spreading area-stiffness relationship and the non-monotonic speed-stiffness relationship, as reported in many experiments [5–7]. Another important addition in our current study compared to our previous model is the inclusion of the cytoskeletal strain-stiffening effect. This effect plays a crucial role in accurately quantifying both cell spreading and migration efficiency of cells with adhesion reinforcement (see Fig. S19). Ignoring this effect could lead to a significant overestimation of the spreading area on stiff substrates, leading to a migration behavior that nearly comes to a halt.

Our simulations on viscoelastic substrates also reveal the influence of substrate viscosity on cell migration by tuning the engagement of the adhesion reinforcement regime. On a substrate with low instantaneous stiffness (*K*_*a*_ + *K*_*e*_ *< K*_0_) where the adhesion reinforcement regime is absent, we have analytically shown the presence of a biphasic traction force-viscosity relationship at an FA site, with the traction force maximized on fast-relaxing substrates 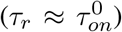. The analytical relation helps us explain the simulation results, which quantitatively agree with the experiments. On substrates with a large instantaneous stiffness (*K*_*a*_ + *K*_*e*_ ≥ *K*_0_), the substrate viscosity can greatly enhance the cell spreading area due to the engagement of the adhesion reinforcement regime. Notably, although the net traction force also increases with viscosity, we observe a biphasic speed-viscosity profile with a very large additional stiffness of *K*_*a*_ ≫ *K*_0_. This is because high viscosity on a very stiff substrate triggers strong adhesion reinforcement, which induces a large cell spreading area, in turn raising the nucleus drag force that counteracts the net traction force.

Finally, we have investigated the influence of substrate viscoelasticity on directed migration. Our model successfully captures the robust durotaxis behavior of the MDA-MB-231 cells with talin-/vinculin-mediated clutch reinforcement on elastic gradient substrates. These results complement our previous findings regarding the talin-low cells that operate without adhesion reinforcement and thus exhibit the co-occurrence of durotaxis and anti-durotaxis on gradient substrates. As a result, our investigations have successfully replicated the complete spectrum of directed migration behaviors displayed by MDA-MB-231 cells reported in Ref. [16]. This underscores the pivotal influence of the adhesion reinforcement regime in governing cell durotactic responses on gradient substrates. Furthermore, our simulated cells also exhibit both viscotaxis and anti-viscotaxis, with all cells migrating towards the fastest-relaxing substrate region where the traction force at an FA site is maximum when 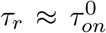. This reveals the critical role of stress relaxation in modulating the migration direction.

Our framework has inherent physical and mathematical limitations in that it is built by coupling cellular processes that are not yet fully understood. For instance, the model only takes into account the active regimes of the actin cytoskeleton at the sub-cellular level using the motor-clutch model. A rigorous modeling of the actin cytoskeleton could involve adopting the active gel theory, which characterizes dynamic structures formed by the interactions of actin filaments and associated proteins [70–72]. Moreover, the interplay between the cytoplasmic flow and the actin cytoskeleton plays a pivotal role in cellular function and behavior [73]. Incorporating fluid mechanics into the active gel theory of the cytoskeleton can enrich our comprehension of the dynamic nature of cells and enable us to understand certain unresolved phenomena, such as the transition between amoeboid and mesenchymal migration [74, 75].

These limitations notwithstanding, our theory provides a valuable tool to understand how the intricate interplay between cellular signaling and mechanics along with cell-ECM interactions regulates migration on viscoelastic substrates. Hence, the theory paves the way for the development of novel experimental platforms for the manipulation of cell migration patterns.

## Supporting Material

Supplementary text, 12 figures, and two tables are available in the appended PDF document.

## Author Contributions

W.S.: conceptualization, formal analysis, investigation, software, validation, visualization, writing – original draft, review and editing; C.N.K.: conceptualization, formal analysis, project administration, resources, validation, writing – original draft, review and editing.

## Acknowledgements

We thank the College of Science at Virginia Tech for financial support.

## Data availability

The source codes of the simulations can be downloaded at https://github.com/nadirkaplan/adhesion-reinforced_migration.

## Appendix

### A1 Rate coefficients in Rho GTPase reaction-diffusion model

Definitions of rates 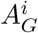 in Eq.15 accommodate autoactivation and antagonistic effects as well as the feedback from the mechanical deformations of the cell membrane. The RhoA activation rate 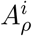 at the *i*^*th*^ vertex is composed of three terms,

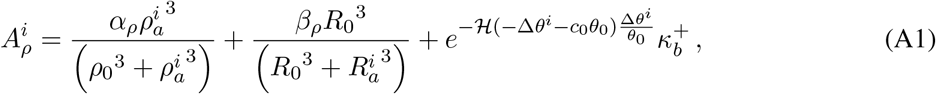

where *R*_0_ and *ρ*_0_ are reference levels of the active Rac1 and RhoA on the vertex, respectively. The first term takes into account positive feedback from the active RhoA itself. The magnitude of the autoactivation effect is represented by *α*_*ρ*_ [76, 77]. The second term describes the mutual inhibition effect between Rac1 and RhoA with a rate *β*_*ρ*_ [76, 77]. The last term couples the active level of RhoA to the mechanical deformations of the cell at the *i*^*th*^ vertex. H() denotes the Heaviside step function. The initial angle at a cell vertex is labeled by *θ*_0_, and Δ*θ*^*i*^ ≡ *θ*^*i*^ − *θ*_0_ denotes the change in angle at the vertex (Fig. 1 E). Here we assume that the active rate of RhoA increases when the vertex angle satisfies Δ*θ*^*i*^ *<* −*c*_0_*θ*_0_ due to the cell polarization.

Similarly, Rac1 activation rate 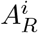 at the *i*^*th*^ vertex also consists of three terms,

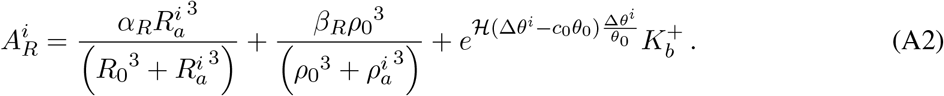

We assume that the active rate of Rac1 soars once the increase of the vertex angle exceeds the threshold *c*_0_*θ*_0_ due to the cell contraction. Coupling Rac1 and RhoA to the mechanical deformations enables recurrent polarization and the recovery of cell deformations. Specifically, an excessive increase in *θ*^*i*^ at the retracting cell rear builds up 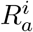 to increase actin polymerization and thus reverses the retraction at the end of each migration step. A large decrease in *θ*^*i*^ at the protruding cell front boosts *ρ*_*a*_ that increases actomyosin contraction to resist further protrusion.

### A2 Analytical derivation of the soft substrate force and deformation

We have derived an analytical expression for the substrate force *F*_*sub*_ at a single FA site to explain the influence of viscosity on a soft substrate. The derivation focuses on isolated motor-clutch dynamics that can also be numerically solved by Eq. 2–6 (Fig. S15 A). For a steady-state solution, we neglect the time-dependence of the variables *N*_*c*_, *N*_*m*_, *F*_*p*_ by setting 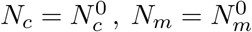, *F*_*p*_ = 0, which are otherwise determined by the chemo-mechanical dynamics at the cellular scale. It is important to note that our goal is not to replicate the substrate force at each FA site of the proposed whole-cell model but to decipher the mechanism of the non-monotonic profile between the viscosity and substrate force on soft substrates. Such motor-clutch analyses that ignore the details of the global dynamics prove to be effective in understanding the influence of matrix mechanics on traction forces [17, 23, 45, 46]. To that end, the analytical derivations presented are aimed at mathematically demonstrating the effect of viscosity.

The molecular clutches bind between the actin filaments and the substrate with a reference association rate 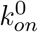, or equivalently at a timescale 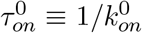. At any instant, the number of the associated clutches is 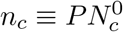. The substrate force builds up at an FA site with the actin filaments moving toward the cell nucleus with the retrograde speed 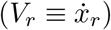. The total restoring force of the bounded clutches is balanced by the substrate deformation. By first considering an elastic substrate, we have *K*_*e*_*x*_*sub*_ = *n*_*c*_*K*_*c*_ (*x*_*r*_ − *x*_*sub*_). Since *n*_*c*_ ≫ 1, it is easy to show that the clutch deformation (*x*_*r*_ − *x*_*sub*_) is insignificant compared with the substrate deformation *x*_*sub*_, or *x*_*r*_ ≈ *x*_*sub*_ [45]. Therefore, we have the force-velocity relation in an alternative form,

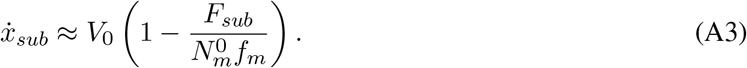

On an elastic substrate, Eq. A3 only contains one unknown *x*_*sub*_ and can be solved with the initial condition *x*_*sub*_|_*t*=0_ = 0,

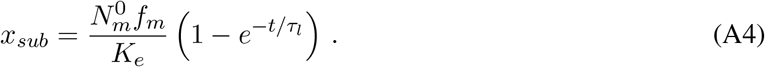

The substrate force is thus given by,

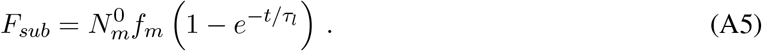

The timescale 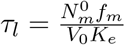 sets the substrate deformation rate and can approximate the lifetime of a binding– unbinding cycle in the FA dynamics [28]. By comparing the substrate deformation timescale *τ*_*l*_ with the binding timescale 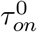, we defined a threshold stiffness 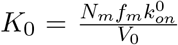 (Table. 2). On soft substrates with *K*_*e*_*/K*_0_ *<* 1, the clutch tension builds up slowly, and a long lifetime 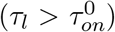 allows for the association of a large number of clutches, i.e., 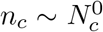. On stiff substrates, the traction force increases rapidly, and a short lifetime 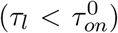 limits the clutch association. Consequently, the force of a single engaged clutch increases significantly (*f > f*_*cr*_ in Eq. 3), triggering adhesion reinforcement.

By describing a viscoelastic substrate by the SLS model, the substrate force-displacement relation is given by

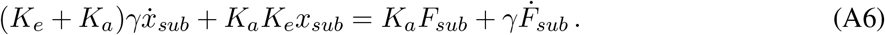

Since Eq. A3 still holds on viscoelastic substrates as long as the instantaneous stiffness is below the threshold, i.e., *K*_*e*_ + *K*_*a*_ *< K*_0_, we substitute Eq. A3 into Eq. A6 to eliminate *F*_*sub*_, which gives

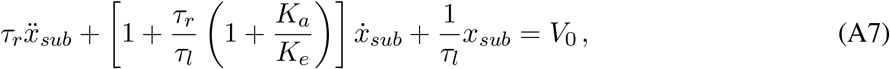

where *τ*_*r*_ ≡ *γ/K*_*a*_ defines the relaxation timescale. With the initial conditions *x*_*sub*_ = 0 and *F*_*sub*_ = 0 at *t* = 0, the solution of the substrate force is obtained as,

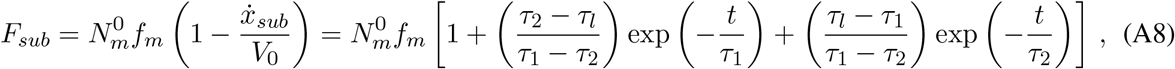

where *τ*_1_ and *τ*_2_ represent two characteristic times with 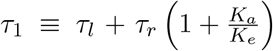, and *τ*_2_ ≈ *τ*_*r*_*τ*_*l*_*/τ*_1_. For soft substrates with a low equilibrium stiffness *K*_*e*_ ≪ *K*_*a*_ (i.e., *K*_*a*_*/K*_*e*_ ≫ 1), we have 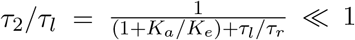 and 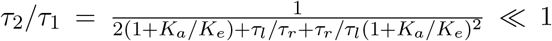. We can thus simplify the equation as

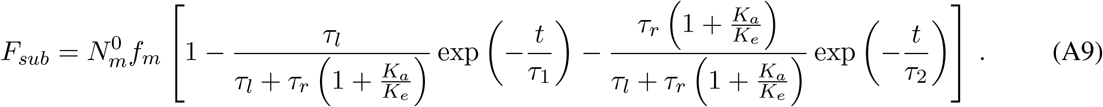

The solutions have good agreement with the numerical results prior to the clutch rupturing at *t* ≈ *τ*_*l*_ (Fig. S15 B). Eq. A9 can be simplified if we focused on two extreme cases. For a substrate with a short relaxation time 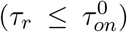, the long duration of the binding-unbinding cycles allows us to neglect the influence of short characteristic time *τ*_2_, which gives,

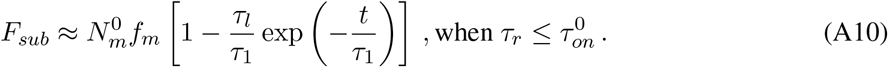

In contrast, a very viscous substrate corresponds to a long relaxation timescale *τ*_*r*_ ≥ *τ*_*l*_, making the contribution of the second term in the square bracket negligible, i.e.,

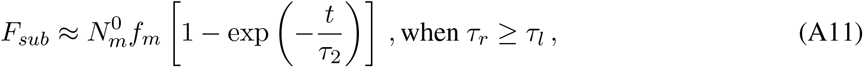

where 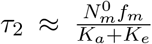, same as the characteristic time on an elastic substrate (Eq. A5) with the stiffness *K*_*a*_ + *K*_*e*_.

With these analytical solutions, we can calculate the average substrate force of binding/unbinding cycle by 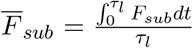. When 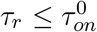, the clutch system preserves the lifetime on an elastic substrate, i.e., 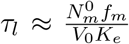. In contrast, a long relaxation time means that the substrate deforms like an elastic substrate with a stiffness *K*_*a*_ + *K*_*e*_. As a result, the shortened lifetime is comparable to the characteristic timescale *τ*_2_ of the substrate deformation when *τ*_*r*_ *> τ*_*l*_. Thus, the average substrate force in two limits is given by,

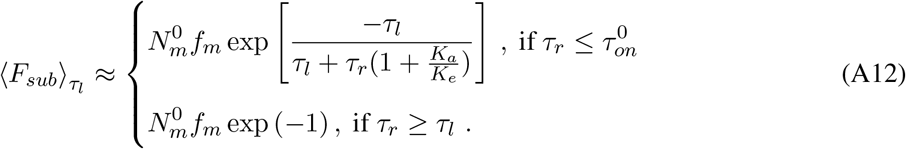

Although the average substrate traction force is difficult to solve for when 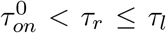, our numerical results demonstrate a fast decrease of the clutch lifetime when 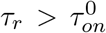, leading to a quick drop of the average substrate force (Fig. S15 C). Taken together, the average traction force must vary in a biphasic fashion with increasing viscosity and achieve a maximum when 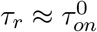 (Fig. S15 D).

The above derivations approximate the dynamics of the traction force at a single FA site with *N*_*c*_ ≈ *N*_*m*_. We next briefly explain the impact of viscosity on the net traction force, which is the traction difference between the cell front and rear. On soft substrates with *K*_*a*_ + *K*_*e*_ *< K*_0_, the traction force decreases with increasing viscosity beyond 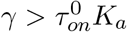 due to the reduced FA lifetime (Fig. S15 C). Consequently, the net traction force must also vary biphasically with increasing viscosity. Due to the regulation of Rho GTPase molecules, the front of the cell has a higher number of clutches than motors 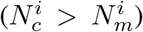, reducing the tension 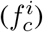 shared by each bond and thus prolonging the clutch lifetime (Fig. S15 E). The extended FA lifetime preserves the high traction force at the front of the cell even when 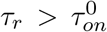 (Fig. S15 F). This explains why the viscosity that maximizes the net traction force is slightly higher than the viscosity that produces the maximum traction force at the rear FA sites.

### A3 Estimation of individual clutch force under adhesion reinforcement

On very stiff substrates, the rapid building of tension 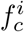 on an individual clutches contributes to a large average bond force far beyond the threshold force (i.e., 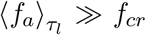), giving rise to very strong adhesion reinforcement effect. The association rate can thus be approximated by 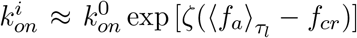. For an engaged clutch with well developed tension 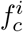, the clutch disassociation rate is mainly dominated by effect of the slip bond, i.e., 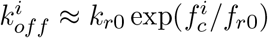. For the saturation of the bounded clutches on very stiff substrates, we must have 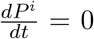 and 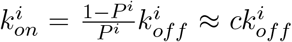 (Eq. 4), where *c* denotes a constant. Therefore, we know that the bond force sustained by an individual clutch can be related to the adhesion reinforcement coefficient by, 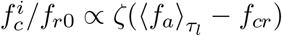. Typically, the maximum of 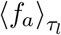 in our numerical analyses is around 10 − 12 *pN*. We can thus further obtain that 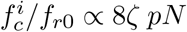.

