## Supporting Information for "A multiscale theory for spreading and migration of adhesion-reinforced mesenchymal cells"

Wenya Shu[1,2] and C. Nadir Kaplan[1,2]*
[1]Department of Physics,
Virginia Polytechnic Institute and State University, Blacksburg, VA 24061, USA.
[2]Center for Soft Matter and Biological Physics,
Virginia Polytechnic Institute and State University, Blacksburg, VA 24061, USA.

### Contents

**S1 Non-negative conditions for retrograde flow speed calculation** **3**

**S2 Implementing the adhesion reinforcement regime in whole-cell model** **3**

**S3 Material parameter fittings** **4**

**S4 Approximation of the drag coefficient** **4**

### S1 Non-negative conditions for retrograde flow speed calculation

The retrograde flow speed at every vertex is calculated by solving Eqs.1-6. Although a maximum of five iterations is enough to achieve convergence for most points (Fig. S1A), computing the minimum retrograde speed typically demands additional iterations (e.g., 20 iterations, Fig. S1B). In our whole-cell model computations, we enforce the $V_r = \max(V_r, 0)$ condition to ensure a non-negative retrograde velocity (Fig. S1C), in case of potential difficulties in achieving the converged solution for the minimum retrograde speed. This prevents the occurrence of negative retrograde velocity, where retrograde actin flows toward the cell periphery—an evident nonphysical behavior.

To further assess the influence of the $V_r = \max(V_r, 0)$ condition on the predictive accuracy of the whole-cell model, we conducted more sophisticated computations that allow a significantly higher maximum iteration count ($\approx 50$) when solving Eqs. 1-6. This enhances the likelihood of achieving a converged solution for the minimum $V_r$ in each loading-unloading focal adhesion cycle. We thus do not enforce the $V_r = \max(V_r, 0)$ condition in sophisticated computation schemes. Our results demonstrate that the predicted cell migration speeds remain almost the same with an efficient scheme that uses fewer maximum iterations but enforces the condition $V_r = max(V_r, 0)$ (Fig. S2).

### S2 Implementing the adhesion reinforcement regime in whole-cell model

The adhesion reinforcement triggered by the talin-unfolding is taken into account by assuming an augmented association rate, denoted as $k_{on}(\langle f_a \rangle_{\tau_l})$. Figure S3 illustrates the calculation of the average clutch force, $\langle f_a \rangle_{\tau_l}$, and its impact on the focal adhesion dynamics. The calculation of the average clutch force, $\langle f_a \rangle_{\tau_l}$, is based on the first binding-unbinding cycle of the focal adhesion dynamics and is given by $\langle f_a \rangle_{\tau_l} = \int_0^{\tau_l} \tilde{f}_c d\tilde{t}/\tau_l$, as shown in Figures S3(B) and S3(D). When the average clutch force, $\langle f_a \rangle_{\tau_l}$, is less than the reinforcement threshold force, $f_{cr} = 2.5\ pN/nm$, the association rate remains constant and equal to $k_{on}^0$. As a result, there is no adhesion reinforcement on soft substrates (Fig S3(B) and S3(C)). However, on stiff substrates ($K_e > K_0 = 5pN/nm$), the single bond force increases rapidly, and only a limited number of bounded clutches form within the short lifetime. This results in a large average clutch force, $\langle f_a \rangle_{\tau_l} > f_{cr} = 2.5\ pN/nm$, as shown in Fig S3(D). An augmented association rate ($k_{on} = k_{on}(\langle f_a \rangle_{\tau_l})$) can accelerate the binding of the unbounded clutches, and the bond force shared by an individual clutch decreases, increasing both the lifetime of the clutch systems and the substrate traction force (Fig S3(E)). On a substrate with very high stiffness ($K_e \gg K_0 = 5\ pN/nm$), the bond force $\tilde{f}_c$ increases rapidly and only a small fraction of the clutches are bounded in the first binding-unbinding cycle, contributing to a large average clutch force, $\langle f_a \rangle_{\tau_l} \gg f_{cr}$. The strong adhesion reinforcement effect can lead to a high binding rate and saturation of the bounded clutches, as illustrated in Fig S3(F).

During the modeling of a whole cell, we need to quantify the effect of the adhesion reinforcement regime at each vertex by determining the average clutch force $\left\langle f_a^i \right\rangle_{\tau_l}$ (Eq 4). The calculation of $\left\langle f_a^i \right\rangle_{\tau_l}$ is based on an isolated motor-clutch model without adhesion reinforcement (Eqs.2-6 with $\zeta = 0$, see the inset in Fig S4). To elaborate, at the time step $t_j$ of the whole-cell model computation, we take the corresponding numbers of clutches $N_c^i$ and motors $N_m^i$, along with the net force resisting protrusion $F_p^i$ of the $i^{th}$ vertex. Next, we apply an isolated motor-clutch simulation with these values of $N_c^i$, $N_m^i$, and $F_p^i$. This simulation is conducted in an independent subroutine and is isolated from the whole-cell modeling process. Its primary objective is to get the dynamics of clutch forces during the initial loading-unloading cycle (see the first cycle in Fig. S3(B)). The mean clutch force is the average clutch force in this first cycle $\left\langle f_a^i \right\rangle_{\tau_l} = \int_0^{\tau_l} \tilde{f}_c^i d\tilde{t}/\tau_l$. The values of $\left\langle f_a^i \right\rangle_{\tau_l}$ obtained from the isolated motor-clutch modeling at every vertex are subsequently transferred to the main routine of the whole-cell modeling (see Fig. S4). Importantly, since variables $N_c^i$, $N_m^i$, and $F_p^i$ change over time in the whole-cell modeling, we must update $\left\langle f_a^i \right\rangle_{\tau_l}$ every $N$ steps of $t_j$. We demonstrate that updating $\left\langle f_a^i \right\rangle_{\tau_l}$ every $N = 4000$ steps can achieve good accuracy while keeping the computational cost reasonable (see Fig. S5).

### S3 Material parameter fittings

Cell stiffness increase due to the strain-stiffening of the cytoskeleton is related to cell spreading area by,

$$\ln\left(\Delta K_{cs} e^{\beta A}\right) = \ln\left(\Delta K_{cs}\right) + \beta A\,, \tag{S1}$$

where parameters $\Delta K_{cs} = 2.8\ pN/\mu m$ and $\beta = 0.00185$ are obtained by the linear fitting of the experimental data in [1] (Fig S6(A)). With these fitting parameters, we calculated the cell spreading areas on elastic substrates and compared them inversely to the experimental results presented in [1] (Fig S6). Clearly, the present model, with $\zeta = 0.5\ /pN$, accurately captures the cell spreading areas in the experiment.

To characterize the viscoelastic substrates made by the interpenetrating networks (IPNs) of alginate, we perform parameter fitting using the Prony series on the representative stress relaxation test data in [2]. The normalized stress in [2] was converted to modulus by considering a constant strain in the stress relaxation test. With the following target equation,

$$\mathcal{E}(t) = \frac{E_\infty + \sum_i E_i e^{-E_i t/\gamma_i}}{E_\infty + \sum_i E_i}\,, \tag{S2}$$

where material parameters $E_i$ and $\gamma_i$ in Prony series are identified in (Table S2) for the fast-, intermediate-, and slow-relaxing substrates, respectively . Due to the limited lifetime of the clutches (Fig S15(C)), the material characterizations are performed by only taking into account the short-term stress relaxation behaviors, i.e., $t \leq 10\ s$. Based on the Prony series coefficients, we defined a constant long-term stiffness $K_e = 0.1\ pN/nm$ and a constant additional stiffness $K_a = 2.5\ pN/nm$ in the standard linear solid (SLS) model (Table S2). The values of the viscosity in the SLS model are defined to approximate the primary relaxation timescale of the IPNs.

### S4 Approximation of the drag coefficient

The drag force acting on the boundary of a migrating cell in 3D matrices is commonly referred to as hydraulic resistance. The drag coefficient is proportional to the fluid viscosity but inversely proportional to the matrix permeability $\kappa_m$ [3],

$$\eta_m = \frac{\mu_c \mathcal{L}_c}{\kappa_m}\mathcal{H}_c \tag{S3}$$

where $\mathcal{L}_c$ and $\mathcal{H}_c$ are characteristic cell length and height, respectively, and both have scales of $1-10\ \mu m$. The viscosity ($\mu_c$) of the extracellular fluid typically has an order of magnitude of $10\ cP$ [4]. The measured hydraulic permeability for the collagen gel is in the range of $\kappa_m = 0.01 - 0.1\ \mu m^2$ [5, 6]. Consequently, the drag coefficient has an effective range of approximately $\eta_m = 1.0 - 100\ pa \cdot s$. In our model, we adopted a value of $\eta_m = 10.0\ Pa \cdot s$. This value is close to the magnitude of the viscous coefficient often used in many 2D models, where the drag force on the cell boundary primarily arises from the viscous responses of the cortex beneath the cell membrane [7, 8]. Our investigation also reveals that the magnitude of the drag coefficient $\eta_m$ has a limited impact on cell migration efficiency (refer to Fig. S7). Nevertheless, it is crucial to highlight that using an excessively small $\eta_m$ should be avoided, as it can lead to numerical instability.

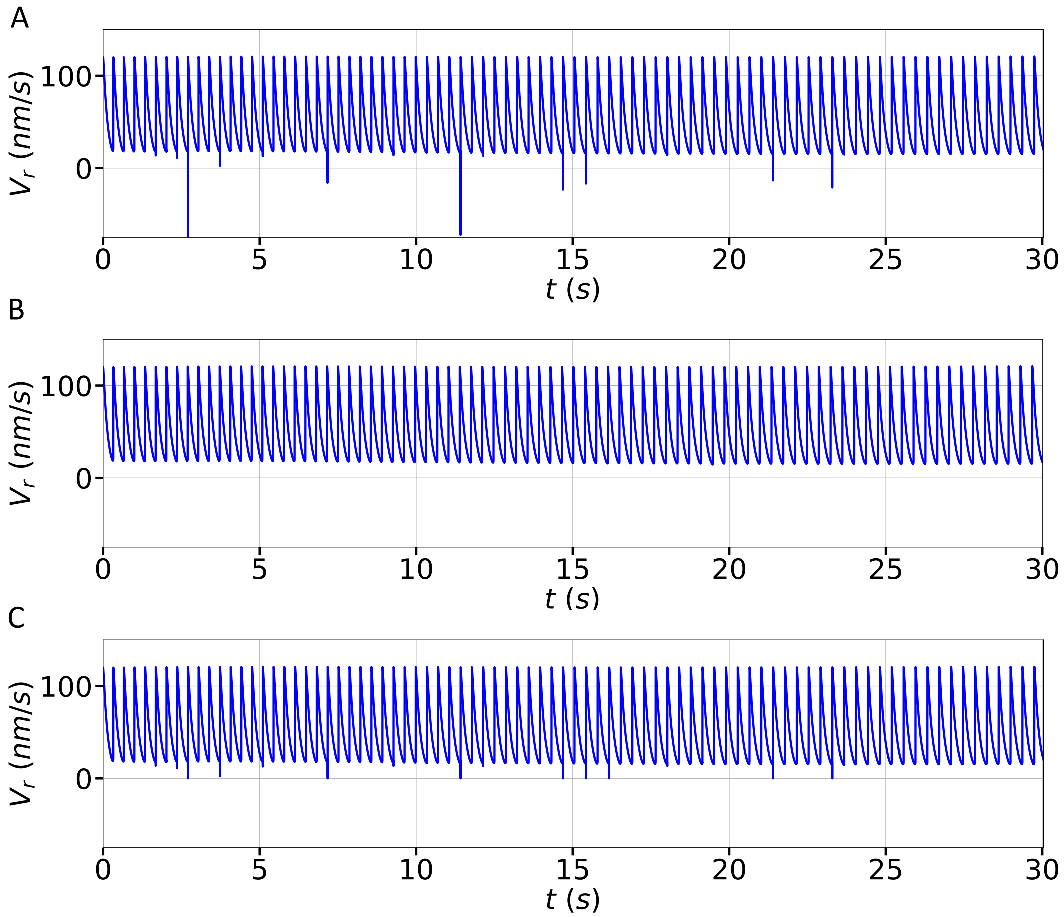

Figure S1: Numerical solutions of the retrograde flow speed at a cell vertex with different numerical treatments. (A) Solutions obtained with a maximum of 5 iterations. (B) Solutions obtained by allowing a maximum of 20 iterations. (C) Solutions corresponding to a maximum of 5 iterations while enforcing the $V_r = \max(V_r, 0)$ condition.

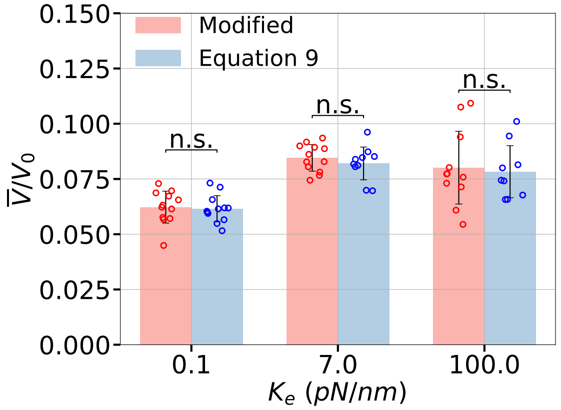

Figure S2: Comparison of average migration speeds between simulations with and without enforcing the condition $V_r = max(V_r, 0)$. Data corresponding to the results from the sophisticated and efficient computations is compared using a t-test, with statistical significance ($p$) denoted as n.s. (not significant) when $p > 0.05$. The scatter plots display all results from simulations ($n \approx 10$), with the standard deviations depicted as error bars.

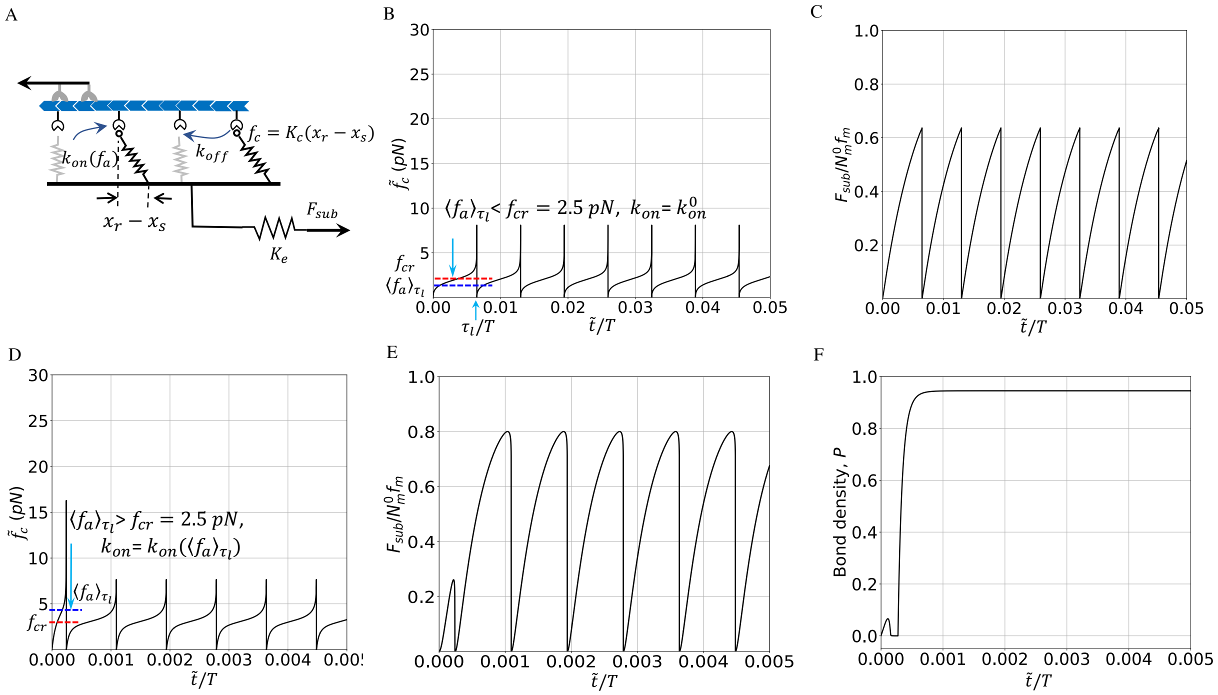

Figure S3: Illustration of the adhesion reinforcement regime with the numerical analysis of an isolated motor-clutch on elastic substrates. (A) Schematic of a motor-clutch model on elastic substrates with stiffness $K_e$. $x_r^i$ and $x_{sub}^i$ denote the displacement of bounded springs and the displacement of the deformed substrate, respectively. $\tilde{f}_c^i$ represents the tension force of an individual spring, while $\tilde{t}$ indicates the simulation time within an isolated motor-clutch modeling context. They are distinct from the variables used in the whole-cell modeling. $k_{on}(\langle f_a^i \rangle_{\tau_l})$ denotes the clutch association rate (Eq 4) and $k_{off}$ is the clutch disassociation rate. (B) The bond force of an individual clutch $f_c$ versus dimensionless time on a soft substrate ($K_e = 1.0\,pN/nm < K_0 = N_m^0 f_m k_{on}^0 / V_0 \approx 5\,pN/nm\,, T$ : migration timescale), which corresponds to a low average bond force $\langle f_a \rangle_{\tau_l} < f_{cr}$. The average bond force (blue dashed line) in the first binding-unbinding cycle is calculated by $\langle f_a \rangle_{\tau_l} = \int_0^{\tau_l} \tilde{f}_c d\tilde{t}/\tau_l$. The threshold force triggering adhesion reinforcement regime $f_{cr} = 2.5\,pN$ is plotted by a red dashed line. (C) Substrate force ($F_{sub}$) as a function of time on soft substrate ($K_e = 1.0\,pN/nm$), which shows that the clutch complex experiences the periodic binding/unbinding dynamics without adhesion reinforcement. (D) Bond force $f_c$ versus time on a stiff substrate ($K_e > K_0$). (E) Substrate force $F_{sub}$ versus time on a stiff substrate ($K_e > K_0$). (F) Strong adhesion reinforcement effect on very stiff substrates $K_e = 50.0\,pN/nm \gg K_0$.

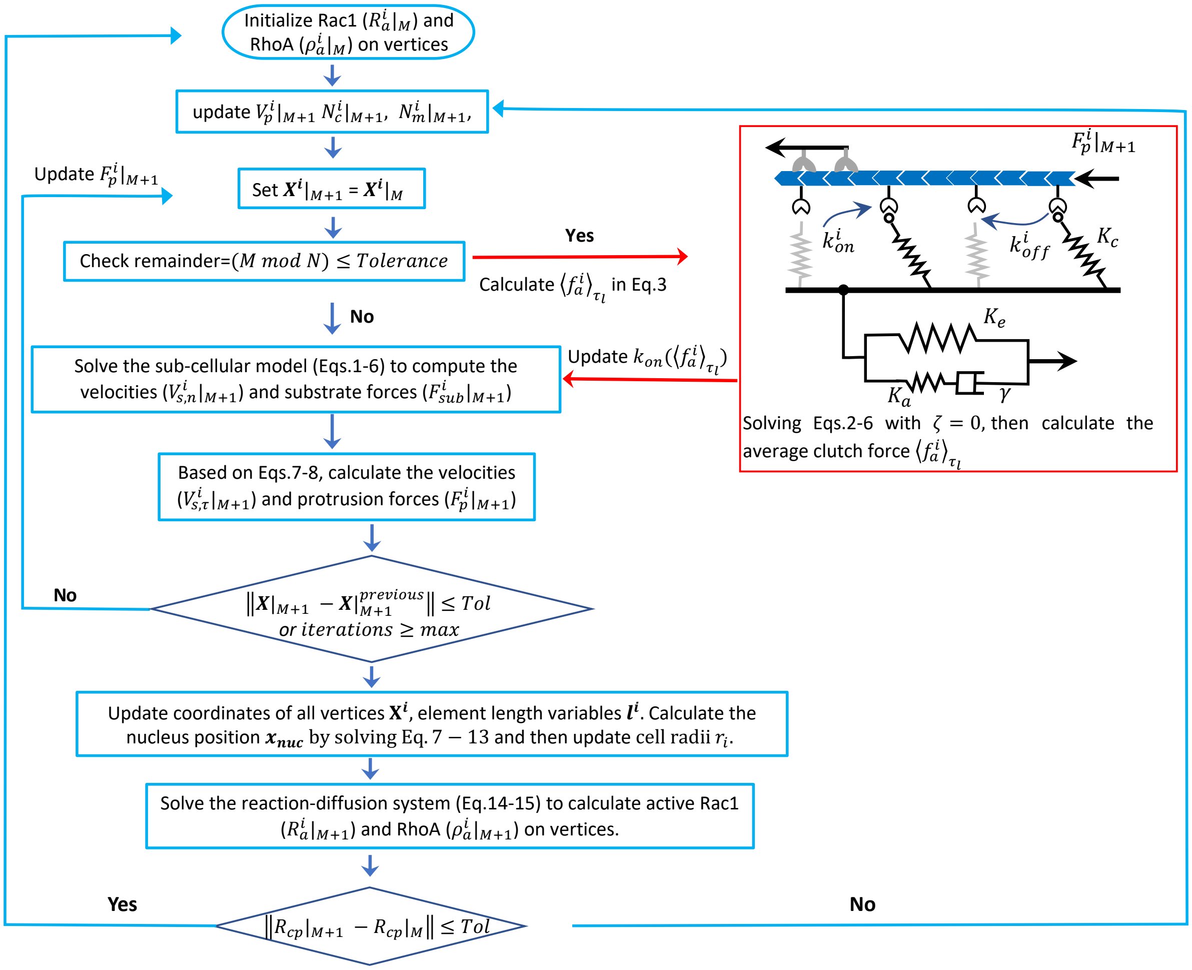

Figure S4: Flowchart of the proposed model implementation. The adhesion reinforcement condition ($\langle f_a^i \rangle_{\tau_l} > f_{cr}$) is checked every $N$ steps. At each time step $(M+1)$, cell radial spreading speed ($V_p^i|_{M+1}$), the number of clutches ($N_c^i|_{M+1}$), and the number of myosin motors ($N_m^i|_{M+1}$) at vertex $i$ are recorded. The position of vertex $i$ at time step $M$ is denoted as $\mathbf{X}^i|_M$. At time step $(M+1)$, radial ($V_{s,n}^i|_{M+1}$) and polar ($V_{s,\tau}^i|_{M+1}$) velocities of vertex $i$ are calculated, as well as the protrusion force ($F_p^i|_{M+1}$) and substrate traction force ($F_{sub}^i|_{M+1}$) of the vertex. A numerical tolerance of $Tol = 10^{-6}$ is set for steady state check.

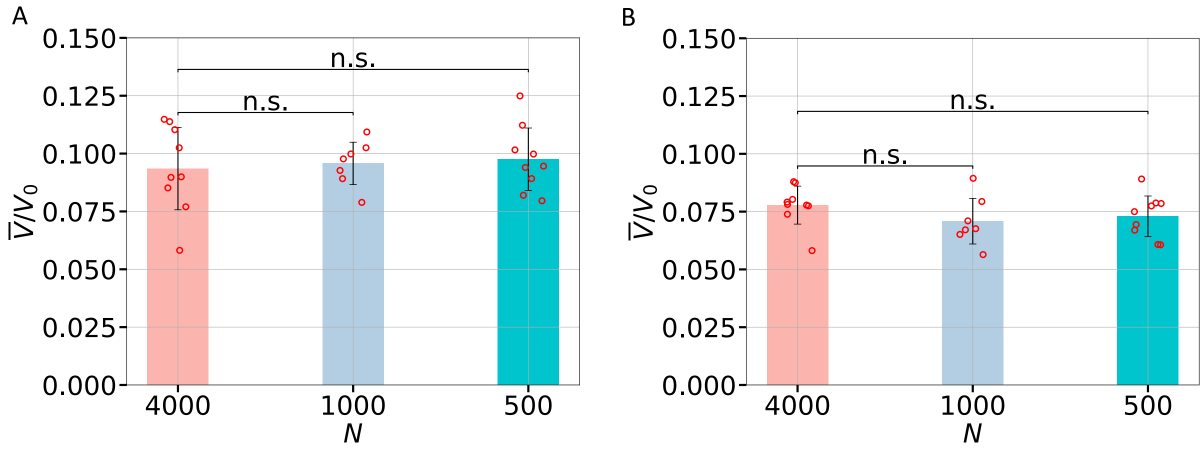

Figure S5: Influence of step interval $N$ for updating $\langle f_a^i \rangle_{|\tau_l}$ on the average speed of cells migrating on stiff substrates with stiffness values of $K_e = 11\ pN/nm$ (A) and $K_e = 100\ pN/nm$ (B).

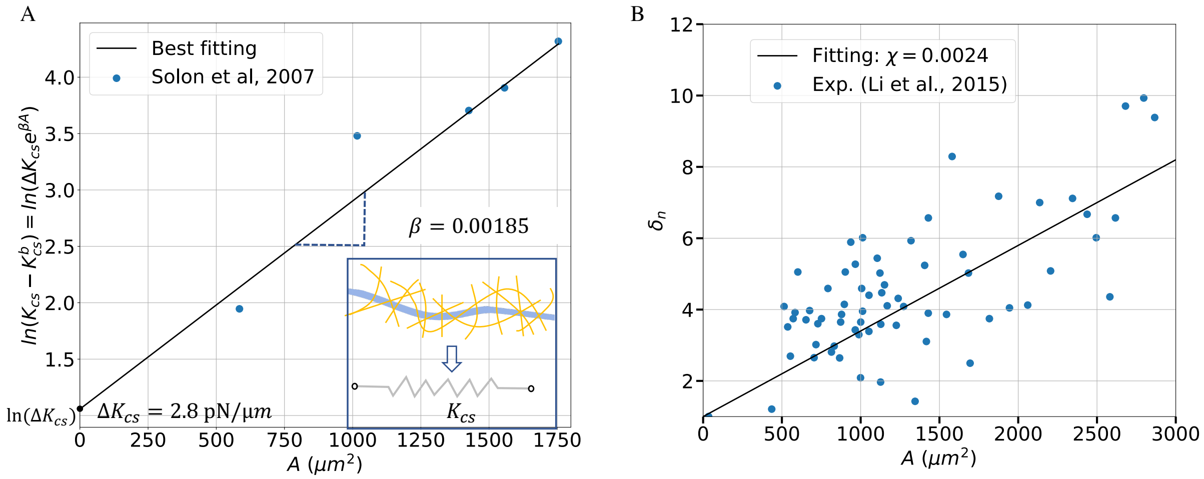

Figure S6: **Primary model parameters obtained from experimental data fitting.** (A) The cytoskeletal biopolymers are represented by the nonlinear elastic elements with the strain-stiffening effect. Parameters in Eq 8 are determined by fitting experimental data [1]. Note that a characteristic intermediate filament length scale of 10 $nm$ is used to convert the modulus ($kPa$) to stiffness ($pN/\mu m$) [9]. (B) Cell nucleus aspect ratio as a function of cell spreading area, with the correlation coefficient $\chi$ in Eq 10 determined by fitting experimental results from [10].

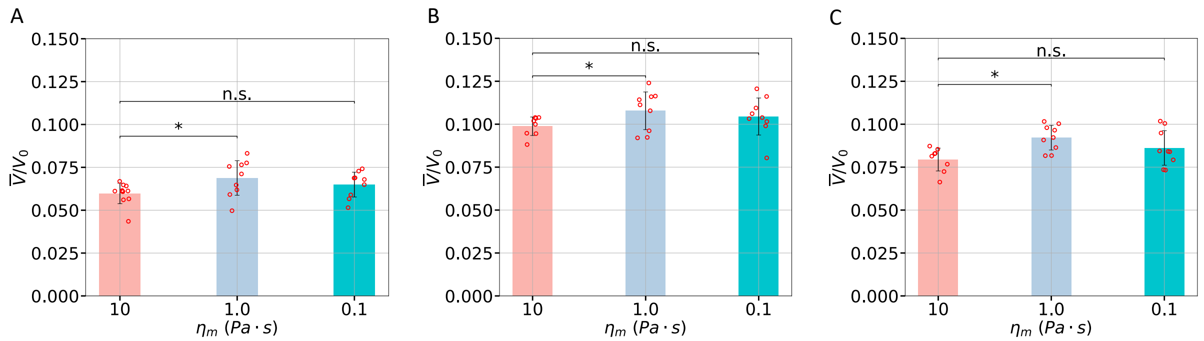

Figure S7: Comparison of average migration speeds obtained from simulations with different drag coefficients ($\eta_m$) as substrate stiffness changes: $K_e = 1.0\ pN/nm$ (A), $K_e = 11.0\ pN/nm$ (B), and $K_e = 100.0\ pN/nm$ (C). Data corresponding to different drag coefficients are compared using t-test, with statistical significance ($p$) denoted as: $*: p < 0.05$, and n.s.: $p > 0.05$. The scatter plots display all results from simulations ($n \approx 10$), with the standard deviations depicted as error bars.

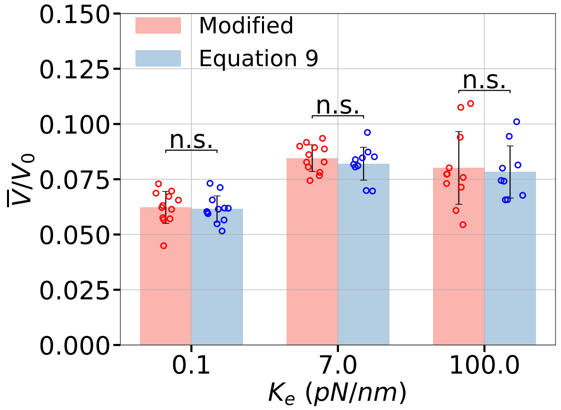

Figure S8: Comparison of average migration speeds between simulations with and without consideration of the relative velocity of the nucleus to the cytoplasm. We use the centroid velocity of the deformed cell shape to represent the cytoplasm velocity around the nucleus. The centroid velocity is calculated by $\dot{\mathbf{x}}_{cp} = (\mathbf{x}_{cp|M+1} - \mathbf{x}_{cp|M})/\Delta t$. The centroid position is calculated as the weighted average of the centroids $\mathbf{c}_i$ of triangles formed by adjacent vertices and the nucleus, with the weights being the areas $A_i$ of these triangles, i.e., $\mathbf{x}_{cp} = \sum_{i=1}^{N} A_i \mathbf{c}_i / \sum_{i=1}^{N} A_i$ . Data corresponding to different forms of Equation (9) is compared using a t-test, with statistical significance ($p$) denoted as n.s. (not significant): $p > 0.05$. The scatter plots display all results from simulations ($n \approx 10$), with the standard deviations depicted as error bars.

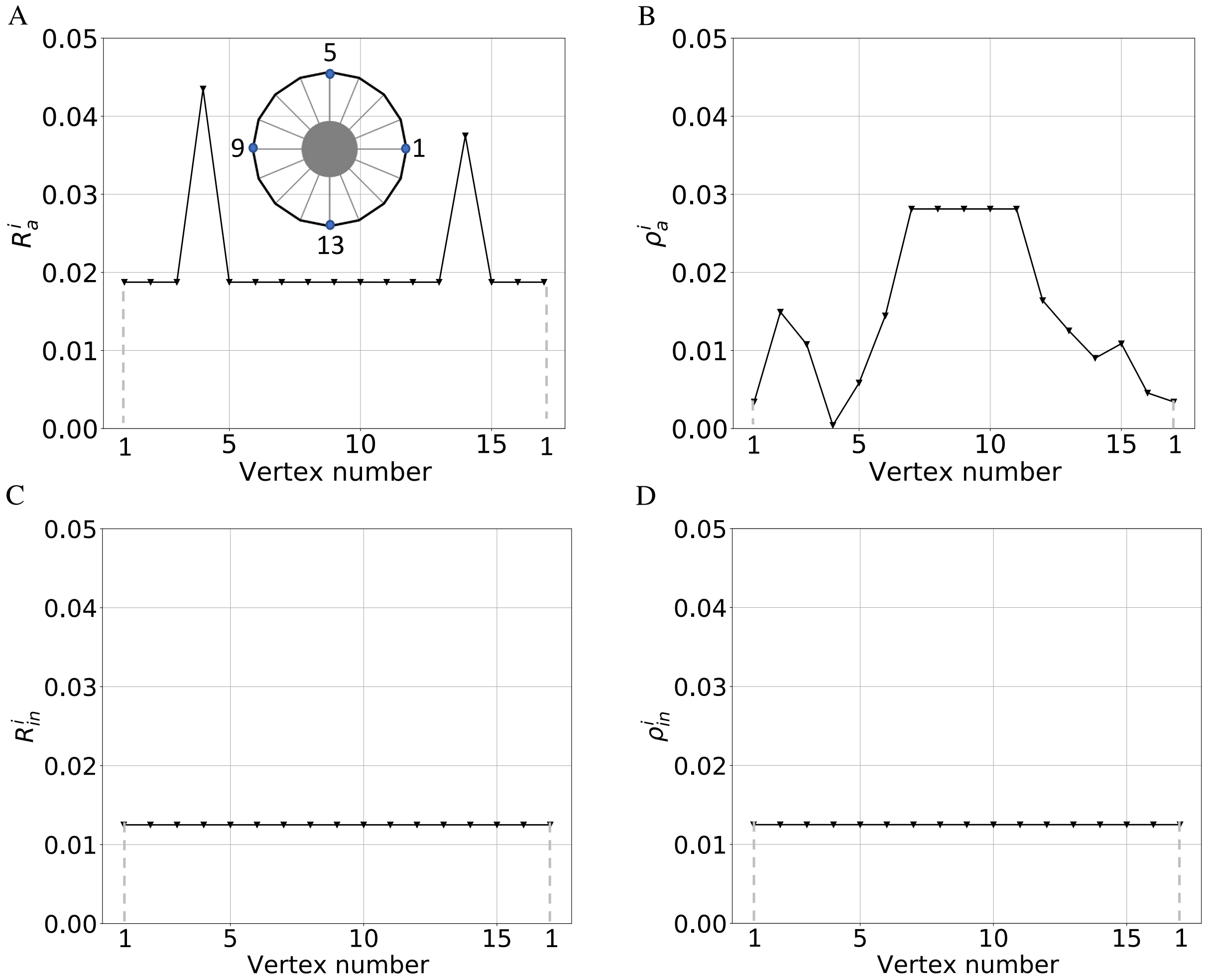

Figure S9: Initial conditions of the membrane-bound Rac1-RhoA signals for the directional migrations. (A) The representative initial condition of the active Rac1 signal $R_a^i$ across all vertices ($N = 16$). Cell shape at $t = 0$ $s$ illustrates the numbering of vertices. (B) Polarized distribution of the active RhoA signal $\rho_a^i$ at $t = 0$ $s$ with the uniform high concentration at the leftmost five vertices. (C-D) All other types of membrane-bound proteins, i.e., $\rho_a^i$, $R_{in}^i$, $\rho_{in}^i$, are uniformly distributed on vertices. Since the total concentration of Rac1 and RhoA are each conserved, the initial cytoplasmic concentration $G_{cp}(t=0)$ , $G \equiv R$ , $\rho$ can be determined by $G_{cp} = 1 - \sum_{i=1}^{N} G_a^i - \sum_{i=1}^{N} G_{in}^i$ .

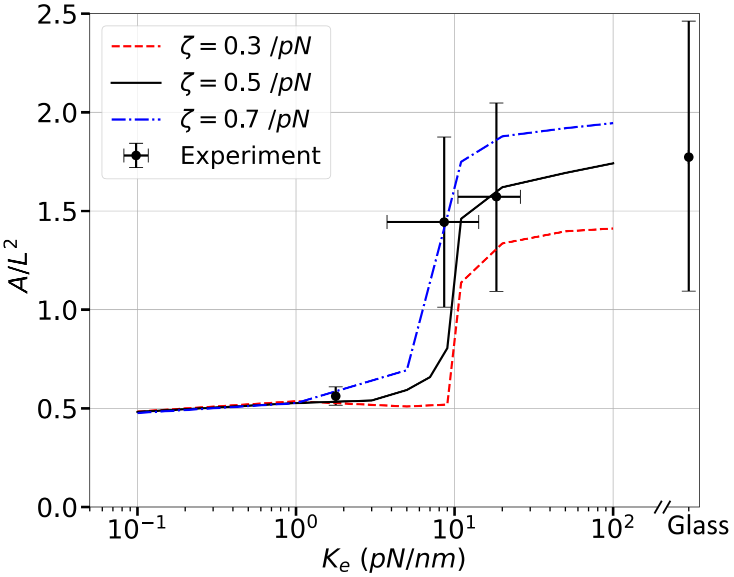

Figure S10: The relationship between the dimensionless cell area and matrix stiffness with different adhesion reinforcement coefficients ($\zeta$ in Eq 5). The spreading area A of a cell is calculated by averaging cell areas over the last $10^6$ steps. The experimental data from [1] is shown as dots with error bars.

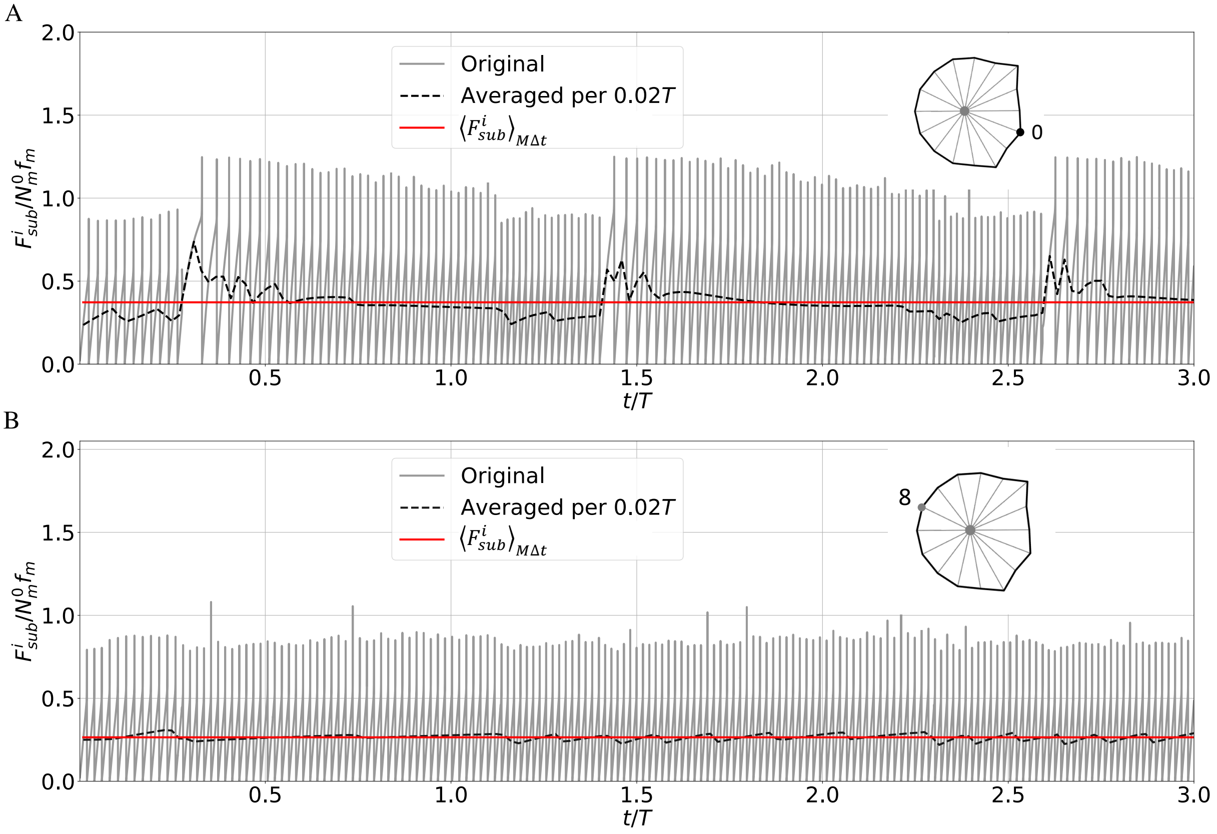

Figure S11: Focal adhesion dynamics at FA sites in the simulations with the proposed whole-cell model. (A) Illustration of binding/unbinding force cycles (full line) at an FA site on the front cell edges and the time-averaged traction force per $0.02T$ (i.e., 4000 steps) versus time (dashed line). (B) Illustration of the focal adhesion dynamics at an FA site on the rear edges of a cell. The red line gives the fitting average traction force $\left\langle F^i_{sub} \right\rangle_{M\Delta t}$ at an FA site, which is reported in Fig 2B.

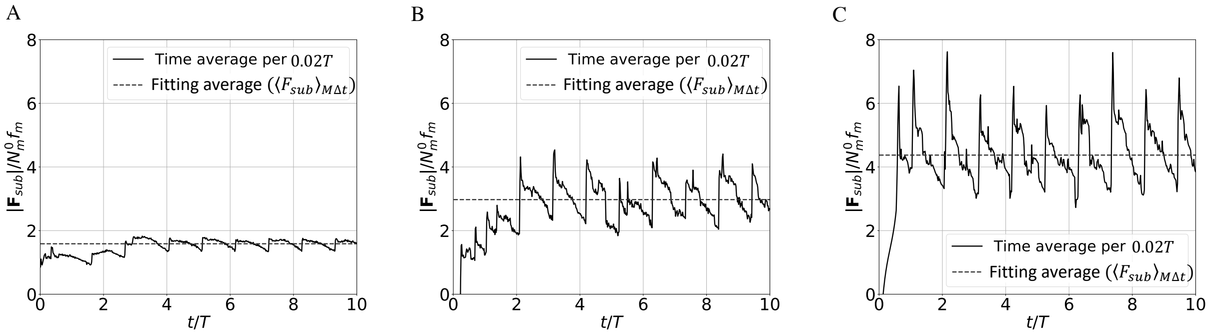

Figure S12: Time-history of the net traction force ($\mathbf{F}_{sub} = \sum_{i=1}^{16} \mathbf{F}^i_{sub}$) and calculation of the average ($\langle \mathbf{F}_{sub} \rangle_{M\Delta t}$, dashed line) for cells on different elastic substrates with stiffness values of (A) $K_e = 2pN/nm$, (B) $K_e = 10pN/nm$, and (C) $K_e = 100\ pN/nm$. The full line represents the time-averaged net traction force per $0.02T$. Note that the magnitude of the vector $\mathbf{F}$ is represented by $|\mathbf{F}|$.

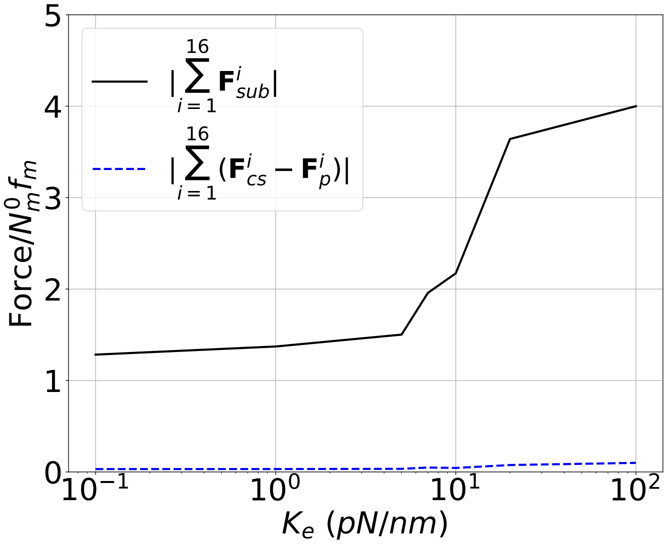

Figure S13: Comparison of net traction force ($\sum_{i=1}^{16} \mathbf{F}_{sub}^i$) and the sum of intracellular forces ($\sum_{i=1}^{16} \mathbf{F}_{cs}^i - \mathbf{F}_p^i$).

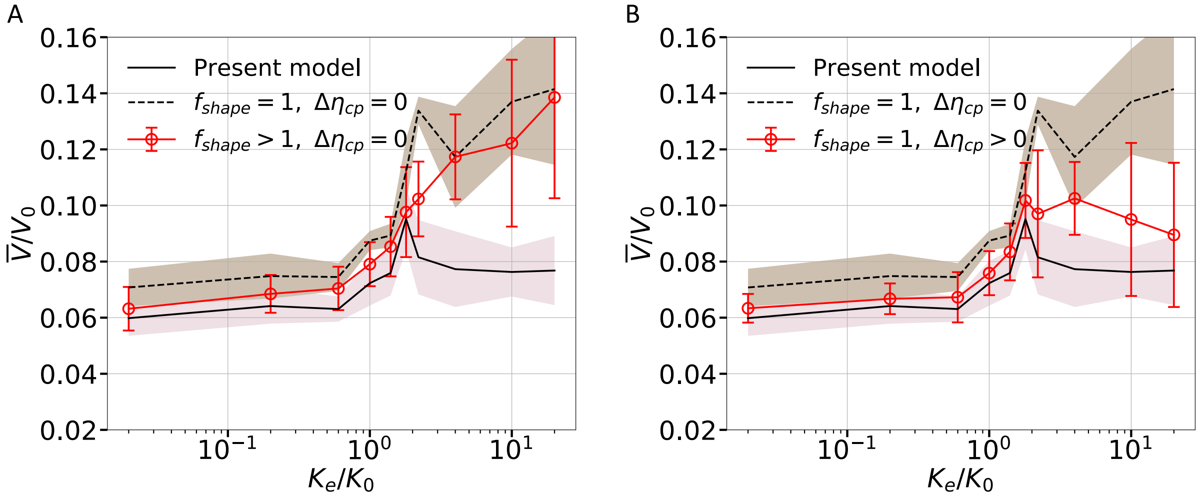

Figure S14: The impact of the shape factor and the cytoplasm viscosity increase on the relationship between the dimensionless mean migration speed $\overline{V}/V_0$ and the dimensionless substrate elastic stiffness $K_e/K_0$ . (A) Influence of shape factor: Comparison between the results from the simulations that consider only the change in the shape factor (dot-line), the full model presented in the manuscript (solid line), and the model assuming a constant viscous drag force on the nucleus (dashed line). (B) Influence of cytoplasm viscosity increase: The results from the simulations that consider only the cytoplasm viscosity increase (dot-line) are compared to the results obtained from the full model presented in the manuscript (solid line) and the model assuming a constant viscous drag force on the nucleus (dashed line).

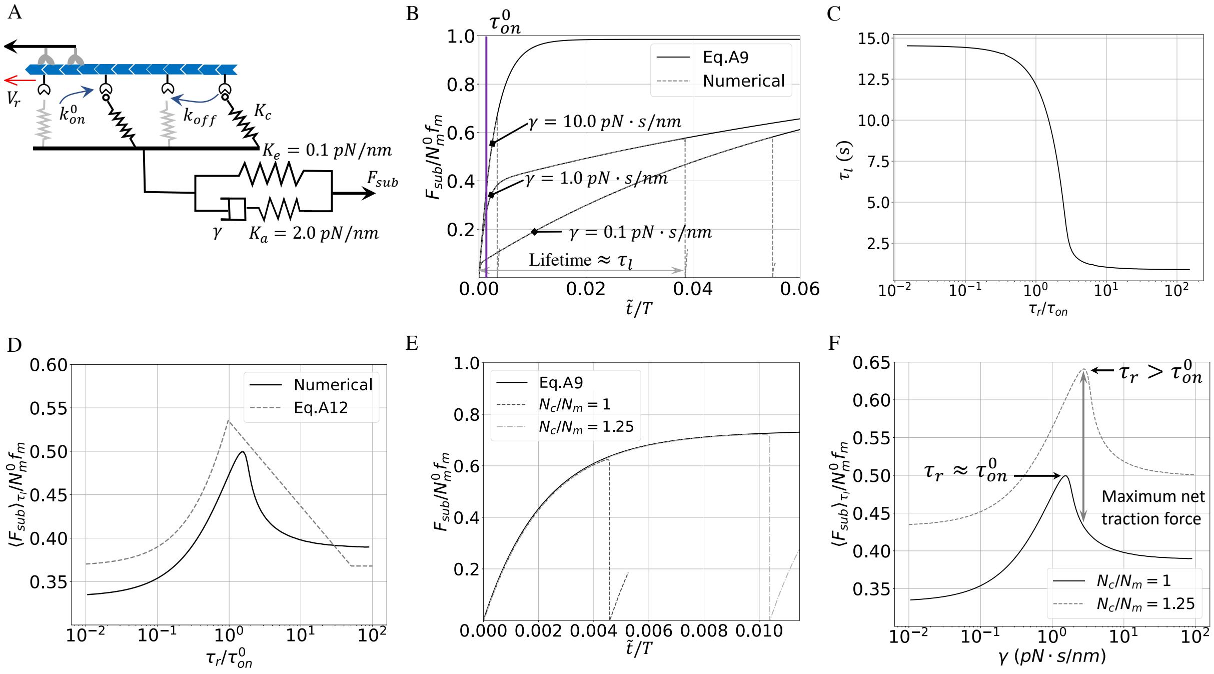

Figure S15: FA analysis investigating mechano-sensing responses to viscosity at the sub-cellular level. (A) An isolated motor-clutch model with no influence from chemical signaling or mechanical forces from the cell body. The adhesion reinforcement mechanism is not considered, as it has an insignificant influence on the soft substrates. (B) Analytical (full) versus numerical (dashed) substrate force dynamics on viscoelastic substrates with different viscosity ($\gamma = 0.1\,, 1.0\,, 10\,pN \cdot s/nm$). (C) The lifetime $\tau_l \approx \tau_l$ of the clutch binding/unbinding cycle versus the ratio of the material timescale to the clutch binding timescale, $(\tau_r/\tau_{on}^0)$. (D) The mean dimensionless substrate force is plotted against $\gamma$. The numerical results, calculated as $\left\langle F_{sub}^i \right\rangle_{\tau_l} = \left( \int_0^{t_f} F_{sub}\, dt \right) / t_f$, are shown as a dashed line. The analytic approximation is shown as a full line. (E) The comparison of dimensionless substrate force dynamics for variations in the clutch number ($N_c = 125$ and $N_c = 100$) is shown. The analytical results are plotted as a full line for comparison. (F) The numerical average substrate force ($\left\langle F_{sub}^i \right\rangle_{\tau_l} / N_m^0 f_m$) is plotted against $\gamma$ for $N_c = 125$ and $N_c = 100$, respectively.

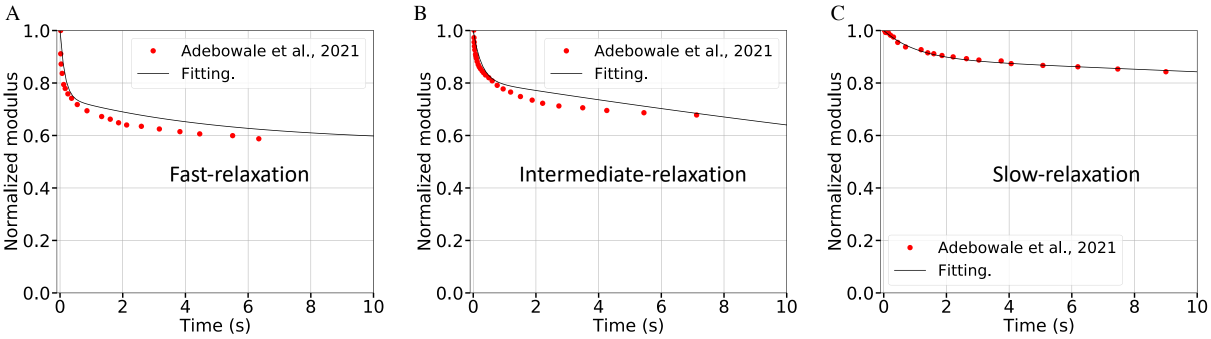

Figure S16: Parameter identifications of the interpenetrating networks (IPNs) of alginate for 2D cell migration studies [2]. Stress relaxation tests of the fast-relaxing (A), the intermediate-relaxing (B), and the slow-relaxing (C) IPNs are fitted by the Prony series (Eq S2.

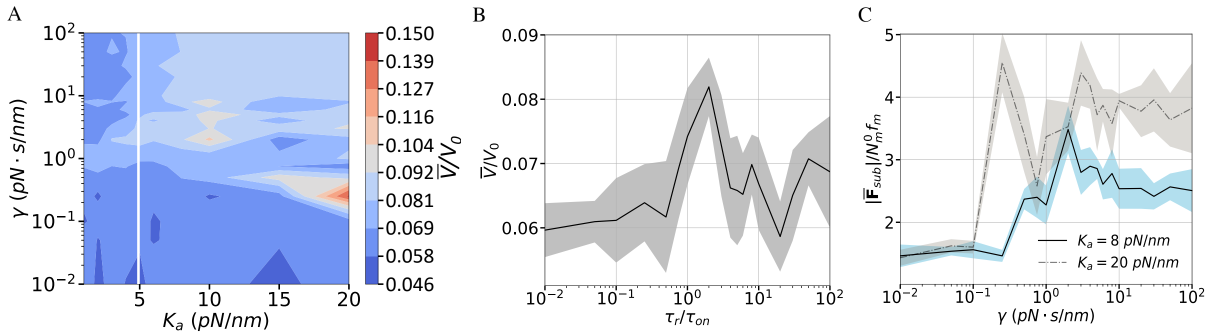

Figure S17: Effects of ECM viscosity on cell migration speed. (A) A contour plot of the dimensionless cell migration speed $\overline{V}/V_0$ as a function of the additional stiffness $K_a$ and substrate viscosity $\gamma$. All simulations use a constant long-term stiffness, $K_e = 0.1\ pN/nm$. The white solid line indicates the threshold stiffness ($K_a + K_e = K_0 = 5\ pN/nm$). The contour plot includes 180 data points with over n = 5 simulations at each point. (B) $\overline{V}/V_0$ and its standard deviation (shaded area) versus the ratio $\tau_r/\tau_{on}^0$ on soft substrates ($K_a = 2\ pN/nm$). (C) The dimensionless mean and standard deviations of the time-averaged net traction force as a function of $\gamma$ on stiff substrates with $K_a = 8\ pN/nm$ and $K_a = 20\ pN/nm$, respectively.

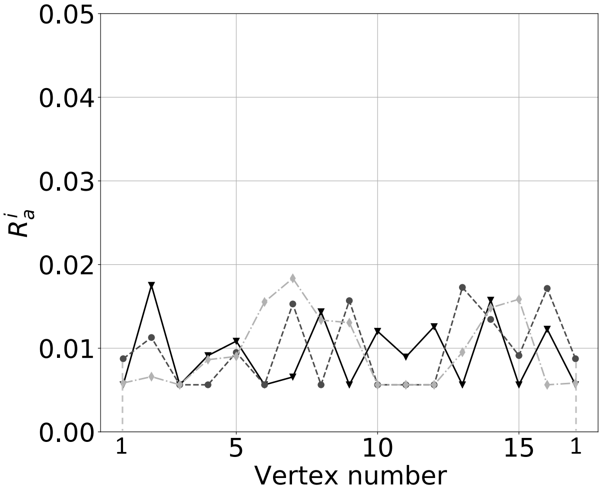

Figure S18: Random distributions of the membrane-bound Rac1 in the active form across all vertices $N = 16$ for migrations ($R_a^i$) on substrates with gradients.

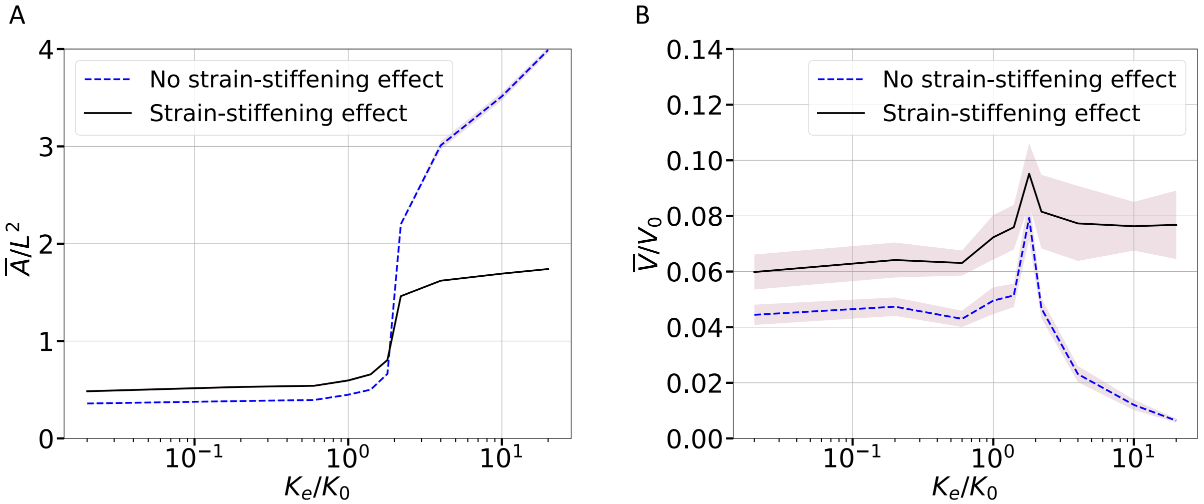

Figure S19: Influence of strain-stiffening effect on cell spreading and migration speed. (A) The dimensionless mean cell area $\overline{A}/L^2$ as a function of the dimensionless substrate elastic stiffness $K_e/K_0$ . (B) The simulated dimensionless mean migration speed $\overline{V}/V_0$ versus $K_e/K_0$. For comparison, the results of new simulations ($n \geq 5$) with a constant cytoskeletal stiffness are compared to the results considering the strain-stiffening effect.

Table S1: Model parameters with fixed values used in the simulations.

| Symbol | Parameters | Values | Sources | Dimensionless |
|---|---|---|---|---|
| $f_m$ | Single myosin motor stall force | 2.0 $pN$ | [11, 12] | 1/100 |
| $K_c$ | Clutch stiffness | 2.0 $pN/nm$ | [11, 13] | $100\pi$ |
| $k^i_{on}$ | Rate constant of clutch association | 3.0 $s^{-1}$ | [12] | $250\pi$ |
| $k_{r0}$ | Clutch unloaded off-rate | 0.25 $s^{-1}$ | [12, 13] | $125\pi/6$ |
| $k_{c0}$ | Clutch unloaded catch-rate | 120 $s^{-1}$ | [12, 13] | $10000\pi$ |
| $f_{c0}$ | Characteristic catch force | 0.5 $pN$ | [12] | 1/400 |
| $f_{r0}$ | Characteristic rupture force | 1.0 $pN$ | [12] | 1/200 |
| $V_0$ | Unloaded retrograde flow velocity | 120.0 $nm/s$ | [11] | 1.0 |
| $V^0_p$ | Characteristic polymerization rate | 120.0 $nm/s$ | [14] | 1.0 |
| $r_0$ | Cell radius | 5.0 $\mu m$ | [14] | $1/2\pi$ |
| $r_{nuc}$ | Nucleus radius | 2.0 $\mu m$ | [7, 15] | $1/10\pi$ |
| $K_m$ | Membrane stiffness | 15.0 $pN/\mu m$ | [16] | $3\pi/4$ |
| $\eta^i_m$ | Viscosity of the surrounding medium | 10.0 $Pa \cdot s$ | Estimated, see Sec. S4 [3–6] | $3\pi/50$ |
| $K^b_{cs}$ | Baseline cytoskeletal stiffness | 1.25 $pN/\mu m$ | [1, 9] | $\pi/16$ |
| $\Delta K_{cs}$ | Stiffness increment | 2.8 $pN/\mu m$ | Fitting parameter (Fig S6) [1] | $7\pi/50$ |
| $\beta$ | Strain-stiffening coefficient | 0.00185 | Fitting parameter (Fig S6) [1] | |
| $\chi$ | Shape correlation coefficient | 0.0024 | Fitting parameter (Fig S6) [10] | |
| $\zeta$ | Adhesion reinforcement coefficient | 0.5 $/pN$ | Adjusted parameter based on [1] ( Fig S10) | 10 |
| $f_{cr}$ | Threshold force | 2.5 $pN$ | Adjusted parameter based on [1] ( Fig S10) | 1/80 |
| $N^0_c$ | Reference molecular clutch number | 100 | [12] | |
| $N^0_m$ | Reference myosin motor number | 100 | [12] | |
| $M^+_R, M^+_\rho$ | Rac1, RhoA membrane association rate | 0.02 $s^{-1}$ | [17] | $5\pi/3$ |
| $M^-_R, M^-_\rho$ | Rac1, RhoA membrane dissociation rate | 0.02 $s^{-1}$ | [17] | $5\pi/3$ |
| $K^+_b$ | Baseline Rac1 activation rate | 0.3 $s^{-1}$ | This article | $25\pi$ |
| $\kappa^+_b$ | Baseline RhoA activation rate | 0.3 $s^{-1}$ | This article | $25\pi$ |
| $K^-$ | Rac1 deactivation rate ($I^i_R = K^-$) | 0.8 $s^{-1}$ | [18, 19] | $2000\pi/3$ |
| $\kappa^-$ | RhoA deactivation rate ($I^i_\rho = \kappa^-$) | 0.8 $s^{-1}$ | [18, 19] | $2000\pi/3$ |
| $R_0$ | Reference level of the active Rac1 | 3/160 | This article | |
| $\rho_0$ | Reference level of the active RhoA | 3/160 | This article | |
| $\theta_0$ | Vertex angle at $t = 0$ | $7\pi/8$ | Model setup | |
| $c_0$ | Ratio of the threshold angle | 0.3 | This article | |
| $\alpha_R$ | Positive feedback rate on Rac1 | 0.3 $s^{-1}$ | [18] | $75\pi/3$ |
| $\alpha_\rho$ | Positive feedback rate on RhoA | 0.3 $s^{-1}$ | [18] | $75\pi/3$ |
| $\beta_R$ | Rate of Rac1 inhibition by RhoA | 0.3 $s^{-1}$ | [20] | $75\pi/3$ |
| $\beta_\rho$ | Rate of RhoA inhibition by Rac1 | 0.3 $s^{-1}$ | [20] | $75\pi/3$ |
| $D$ | Diffusivity on the membrane | 0.01 $\mu m^2/s$ | [18, 19] | $1/120\pi$ |
| $\eta^0_{cp}$ | Reference cytoplasm viscosity | 860 $Pa \cdot s$ | [21] | $5.76\pi$ |
| $\Delta\eta_{cp}$ | Cytoplasm viscosity increment | 36 $Pa \cdot s$ | This article | $24\pi/125$ |

Table S2: Stress relaxation test data [2] fitted by the Prony series.

| Symbol | Prony series coefficients | SLS model parameters |
|---|---|---|
| Fast-relaxation | $E_\infty = 0.1\,kPa$<br>$E_1 = 0.38\,kPa\,, E_2 = 0.66\,kPa, E_3 = 1.44\,kPa$<br>$\gamma_1 = 1.8\,kPa \cdot s\,, \gamma_2 = 0.1\,kPa \cdot s\,, \gamma_3 = 534\,kPa \cdot s$ | $K_e = 0.1\,pN/nm$<br>$K_a = 2.5\,pN/nm$<br>$\gamma = 1.0\,pN \cdot s/nm$ |
| Intermediate-relaxation | $E_\infty = 0.1\,kPa$<br>$E_1 = 2.0\,kPa\,, E_2 = 0.50\,kPa$<br>$\gamma_1 = 80.9\,kPa \cdot s\,, \gamma_2 = 0.13\,kPa \cdot s$ | $K_e = 0.1\,pN/nm$<br>$K_a = 2.5\,pN/nm$<br>$\gamma = 60.0\,pN \cdot s/nm$ |
| Slow-relaxation | $E_\infty = 0.1\,kPa$<br>$E_1 = 2.2\,kPa\,, E_2 = 0.30\,kPa$<br>$\gamma_1 = 370.0\,kPa \cdot s\,, \gamma_2 = 0.30\,kPa \cdot s$ | $K_e = 0.1\,pN/nm$<br>$K_a = 2.5\,pN/nm$<br>$\gamma = 360.0\,pN \cdot s/nm$ |
